# A meta-analysis resolves the huntingtin interactome into coactivator losses and a robust proteostatic and synaptic gain network

**DOI:** 10.64898/2026.07.06.736704

**Authors:** Manuel Seefelder

## Abstract

Transcriptional dysregulation and proteostatic collapse are cardinal yet mechanistically separate features of Huntington disease (HD), and how the polyglutamine (polyQ) expansion in huntingtin (HTT) rewires its interactome to produce both remains unresolved. We integrated four published HTT affinity-proteomics datasets and contrasted wild-type and polyQ-expanded HTT within one Bayesian model (BayesInteractomics). Of 4,338 proteins, 275 were condition-dependent: the expansion strips HTT of the transcription-activation machinery (Mediator, the ASCOM H3K4-methyltransferase, CREBBP, CDK9) while gaining contacts with the 26S proteasome, HSP70 chaperones and a synaptic and actin-cytoskeletal network, around an intact chaperonin–HAP40 core. This picture emerges only from integration: the datasets overlap so little that a “reproducible in ≥ 2 studies” consensus would recover just ∼ 21% of the high-confidence interactors. By reconciling HD’s transcriptional and proteotoxic arms within one quantitative interactome, this loss-plus-gain model recasts two historically separate disease mechanisms as complementary and nominates prioritised interfaces (HTT–Mediator/ASCOM, HTT–proteasome) for validation and therapeutic targeting.

## Introduction

Huntington disease (HD) is an autosomal-dominant neurodegenerative disorder caused by a CAG-repeat expansion in exon 1 of *HTT*, which lengthens an N-terminal polyglutamine (polyQ) tract in huntingtin (HTT) beyond a pathogenic threshold of 36 residues^1^. HTT is a 350-kDa HEAT-repeat solenoid with no intrinsic catalytic activity; its cellular roles are executed almost entirely through protein–protein interactions, so the consequences of the polyQ expansion are most naturally framed as a *remodelling of the HTT interaction network*^2,3^. Wild-type HTT (wtHTT) — and mutant HTT alike — folds with HAP40 into a stable heterodimer^4,5^. HTT itself scaf-folds dynein/dynactin-dependent transport^6^ and supports BDNF trafficking^7^ — functions documented for huntingtin rather than for HAP40 or the heterodimer. HAP40 occupancy is not incidental to this biology: HAP40 protein levels are huntingtin-dependent, so the two abundances are coupled and both decline in HD^8^ — HTT stabilises HAP40, though a reciprocal interdependence has also been reported^9^. This dependency has recently been formalised as a stoichiometric HTT–HAP40 “rheostat” in which the level of the shared scaffold, rather than HTT alone, tunes its downstream activities^10,11^. How the polyQ expansion remodels the interactome of this constitutive scaffold is therefore central to interpreting the network changes of HD.

Two dominant but historically separate mechanistic threads connect this biology to pathology. The first is transcriptional: expanded polyQ binds and sequesters the acetyltransferase coactivator CREB-binding protein (CBP/CREBBP) and disrupts coactivator-dependent transcription^12,13^, while the activating chromatin mark H3K4me3 is redistributed in HD brain in a manner that modifies disease phenotypes^14,15^. The second is proteostatic: mHTT perturbs the ubiquitin–proteasome system^16,17^ and is routed through selective autophagy^18^, in a cellular context of early, striatum-biased bioenergetic failure^19–21^. Affinity-purification mass spectrometry (AP-MS) has catalogued hundreds of candidate HTT partners^3,22–24^, many of them proposed as mutant-HTT–preferential and advanced as candidate disease mechanisms or drug targets^12,13,16^, yet individual studies differ in species, tag, polyQ length and statistical treatment, are seldom reproduced protein-by-protein, and rarely contrast wtHTT and mHTT inside one combined probabilistic model.

Here we integrate four published huntingtin affinity-proteomics datasets and analyse wild-type (wtHTT) and mutant (mHTT) baits jointly with BayesInteractomics, an open-source Bayesian differential-interactomics framework (outlined below and detailed in the companion methods paper^25^). We restricted the analysis to datasets that satisfied two inclusion criteria — at least three biological replicates per condition, and matched negative controls processed within the same dataset — so that detection consistency and enrichment could be estimated study-internally before any cross-study integration. The four eligible datasets were chosen to span complementary species, tissues and capture chemistries: *(i)* Greco *et al*.^22^ — FLAG–HTT immunoaffinity purification–MS from Q20 and Q140 knock-in **mouse striatum** at 2 and 10 months, integrating label-free and metabolic-labelling quantification (12 HTT and 12 IgG-control IPs); *(ii)* Justice *et al*.^23^ — multi-epitope immunocapture with two epitope-distinct antibody pairs (N-terminal 2B7/4C9 and central 3E10/4E10) across **mouse striatum, cortex and cerebellum** (Q20 vs Q140; 8 and 40 weeks; 81 epitope-resolved IPs against IgG controls); *(iii)* Sap *et al*.^24^ — chemical-**crosslinking** FLAG co-immunoprecipitation of full-length wtHTT and mHTT from **mouse cortex** (control, Q20 and Q140; four replicates each); and *(iv)* Gutiérrez-García *et al*.^26^ — endogenous total-HTT co-immunoprecipitation with label-free MS in **human iPSCderived striatal neurons** (wild-type condition; 3 HTT vs 3 GFP-control IPs). BayesInteractomics^25^ is an open-source Bayesian meta-analytic framework that combines a sequence- and network-based prior with the AP–MS evidence by Bayes’ rule. The prior is the calibrated output of a meta-learner over fourteen features — a deep-learning predictor of direct interaction (a neural network trained on Protein Data Bank contacts from sequence and cross-species network embeddings), STRING functional-association scores and Pfam domain–domain features — so that no single predictor, the neural network included, sets the prior alone; the evidence model fuses three complementary lines of AP–MS evidence (detection consistency, quantitative enrichment and bait dose–response) across the pooled datasets through Bayesian model averaging. For every protein the two are combined into a posterior probability of interaction in each condition, controlled at a Bayesian false-discovery rate, together with a decision-theoretically optimal interaction class. The method, and its validation against simulation ground truth and head-to-head benchmarking against established tools, are described in detail in the companion methods paper^25^. We use the resulting differential map to ask which physiological contacts the expansion *removes* and which aberrant contacts it *creates* — a combined, cross-study wtHTT-versus-mHTT differential re-analysis that extends the integrative direction the most comprehensive of these studies^23^ advocated.

## Results

We describe a protein as *preferential* for a bait when it is a credible interactor of *both* baits but significantly stronger for one, and as *specific* when it is a credible interactor of only one bait. The two arms of the comparison are then the *wtHTT-preferential* set (proteins preferential or specific for wtHTT, i.e. lost on expansion) and the *mHTT-gained* set (proteins preferential or specific for mHTT). The direction of each change is set by the decision-theoretic optimal call (the Bayes action under the full posterior); the differential z-score (Δz = z_wtHTT_ − z_mHTT_) is reported as the accompanying effect-size axis.

### A focused, bidirectional remodelling of the HTT interactome

Across 4,338 proteins evaluated in both conditions, 2,864 received a confident differential call; the overwhelming majority of these were non-interactors of either bait (2,295 both-negative) or were retained without a credible change (294 unchanged). A focused set of 275 proteins received a condition-dependent call (Figure 1). The remaining 1,474 evaluated proteins had replicate coverage too sparse in at least one condition for a confident differential call and were left unclassified. Consistent with mHTT acquiring an aberrant interaction surface, the expansion produced substantially more gains than losses: 205 proteins preferentially engaged mHTT (197 mHTT-preferential plus 8 mHTT-specific) while 70 preferentially engaged wtHTT (50 wtHTT-preferential plus 20 wtHTT-specific). This direction mirrors the underlying single-condition models, in which mHTT recovered more high-confidence interactors than wtHTT (297 vs 196 at BFDR ≤ 0.05).

**Figure 1.**
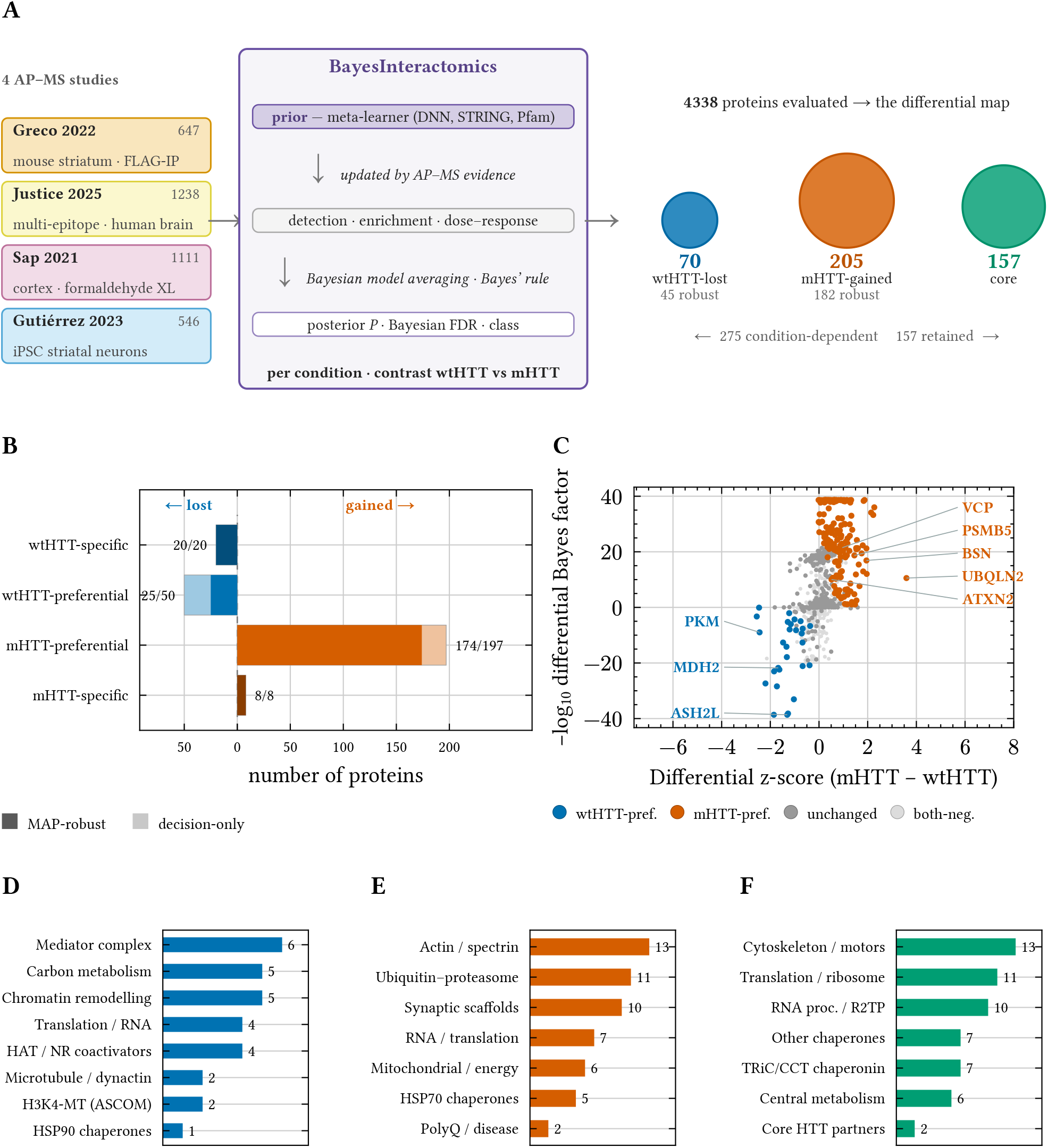
Bidirectional remodelling of the HTT interactome across four pooled AP–MS studies. **(A)** Integration workflow: four heterogeneous wtHTT/mHTT AP–MS datasets (interactor counts shown) are analysed by BayesInteractomics, which combines a *calibrated prior* — the output of a meta-learner over sequence- and network-derived features (a deep-learning PPI predictor being one of them, alongside STRING and Pfam domain features) — with the AP–MS evidence (detection, enrichment and dose– response streams under Bayesian model averaging) by Bayes’ rule, returning per protein and condition a posterior interaction probability, a Bayesian FDR and a decision-theoretic class. Of 4,338 proteins evaluated in both conditions, 275 are condition-dependent (70 wtHTT-lost, 205 mHTT-gained; MAP-robust subsets 45 and 182) alongside a 157-protein constitutive core; circle area is proportional to protein count. **(B)** Differential composition by class, losses left and gains right of zero; solid = calls also recovered by the *maximum-a-posteriori* point classifier (high-confidence), pale = decision-rule only; labels give robust/total. **(C)** Differential Bayes-factor volcano: differential z-score (mHTT − wtHTT) versus −log_10_ differential Bayes factor, coloured by class, with representative hits labelled. **(D–F)** Functional signatures as curated complex/module membership (interactor count per category), bars coloured by class, for **(D)** the wtHTT-lost, **(E)** the mHTT-gained and **(F)** the constitutive-core sets. Genome-background over-representation analysis is given in Supplementary Figure 2 and the per-protein differential map in Supplementary Figure 1.

Both arms were statistically well supported. As a conservative confidence check we retained only proteins whose decision-theoretic call was also recovered by the *maximum-a-posteriori* (MAP) point classification (the MAP-robust tier). The larger mHTT-gained set was the more robust: 182 of the 205 gains were MAP-robust (174 of 197 mHTT-preferential and all 8 mHTT-specific), against 45 of the 70 losses (25 of 50 wtHTT-preferential and all 20 wtHTT-specific; Supplementary Table 2). Interactome-wide, the MAP classification assigned 351 proteins to the mHTT-gained direction and 46 to the wtHTT-lost direction. The mHTT-gained calls therefore carry slightly more point-estimate support than the losses, at a median differential false-discovery rate of effectively zero (< 10^−6^ in both arms; Supplementary Table 2). The robust tiers contain every principal functional module of both arms — on the loss side the Mediator complex, the AS-COM H3K4-methyltransferase (`ASH2L`, `RBBP5`), `CREBBP`, `CDK9` and the glycolytic/chaperone sets; on the gain side the 26S proteasome/ubiquitin machinery, HSP70 chaperones and the synaptic and actin-cytoskeletal network. The expansion therefore drives a focused, *bidirectional* remodelling in which both the discrete coactivator losses and the larger proteostatic and synaptic gain network are individually well supported. We report the decision-theoretic calls as the primary analysis and the MAP-robust subset as a high-confidence tier.

This contrast is displayed as a differential Bayes-factor volcano (Figure 1C); a clustered, per-protein map of the wtHTT and mHTT fold changes across all 275 condition-dependent proteins is given in Supplementary Figure 1.

### The differential signatures are supported across the source datasets

Because the integrated call pools four heterogeneous studies, we asked — before interpreting the signatures biologically — how robustly each differential call is supported across the contributing datasets (Figure 2). Integration is essential here, not cosmetic: because the four studies overlap so little, 79% of the high-confidence (*P* > 0.95) interactors are supported by at most one study individually, so a conventional “reproducible in ≥ 2 studies” consensus would recover only ∼ 21% of them (Supplementary Figure 9). Yet where two or more studies co-detect a protein, the per-study effect magnitudes are statistically homogeneous rather than contradictory (Cochran’s *Q*/*I*^2^: median *I*^2^ = 0, no significant heterogeneity for ∼ 95% of calls and all of the losses; Supplementary Table 3) — the residual scatter reflects under-powering, not disagreement. The per-dataset enrichment volcanoes (Figure 2) confirm that the proteins assigned to each differential class are enriched over control in the datasets that detect them, rather than being artefacts of a single study; the same panels overlay the two directly-profiled HAP40 baits, in which a subset of the wtHTT- and mHTT-class proteins are also enriched, consistent with the constitutive HTT–HAP40 core. We summarise the entire condition-dependent interactome as a single dataset-resolved map (Figure 4), in which every member is coloured by its differential class and annotated with the datasets that support it.

**Figure 2.**
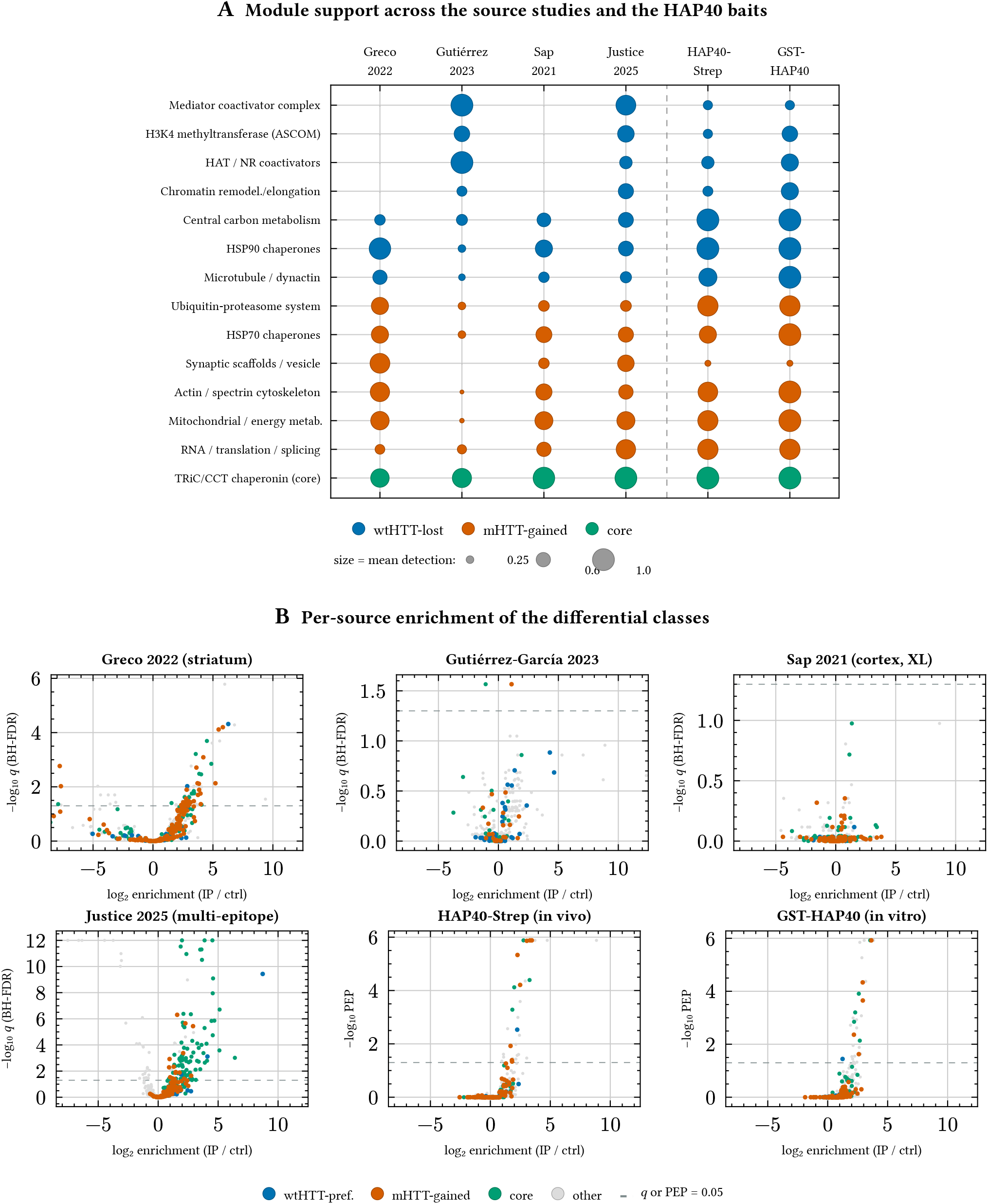
Per-source support of the differential signatures, including the HAP40 baits. **(A)** Source × module map: rows are curated functional modules, columns the four meta source datasets plus the two directly-profiled HAP40 baits (HAP40-Strep, GST-HAP40). Dot radius ∝ module members detected (source studies, raw detection ≥ 50% of replicates; HAP40 baits, *is-detected* flag); fill colour = module direction (blue, wtHTT-lost; vermillion, mHTT-gained; green, constitutive core); shade (pale→solid) = mean per-member detection. The TRiC/CCT core is present in all six. **(B)** Per-source enrichment volcanoes: log_2_ enrichment (bait IP vs control) versus significance — −log_10_ *q* for the meta studies (Welch *p*, BH-corrected, MNAR-imputed), −log_10_ posterior error probability (PEP) for the HAP40 baits’ single-bait BayesInteractomics models — every detected protein coloured by its integrated differential class. Many wtHTT- and mHTT-class proteins are also enriched in the HAP40 baits.

A true model-refit leave-one-study-out test confirmed that this support is not driven by any single dataset. Refitting the entire pipeline four times — each time dropping one study’s protocol(s) and re-running both single-condition models and the differential — left the *direction* of the calls essentially invariant: 269 of the 275 condition-dependent calls (97.8%) kept their direction across all four refits, and at most three reversed in any single refit (`CORO2A`, `DCLK2`, `GRN`, `PSMC1`, `TPPP`, `WDR7` Supplementary Figure 11). What changes under study removal is statistical power, not direction — rather than reversing, calls that lose support collapse to non-significant. This is mild for the single-protocol datasets (40–81% of losses and 34– 49% of gains retained) but pronounced for the dominant Justice *et al*. 2025 dataset, which contributes roughly half the mutant-condition samples and whose removal retains only 6% of the mHTT-gained calls; the mutant arm is accordingly the more power-sensitive of the two. The full-data calls are therefore directionally robust to study removal yet require the pooled evidence to reach significance, consistent with the per-study under-powering documented above (Supplementary Table 3).

### wtHTT preferentially engages the transcription-activation machinery

The proteins lost on expansion describe a strikingly coherent transcription-activation module (Figure 1D; Supplementary Table 4), spanning three layers of the gene-activation machinery. The *Mediator* coactivator complex is the most prominent: six tail- and head-module subunits are wtHTT-specific (Figure 3A; `MED8`, `MED14`, `MED15`, `MED20`, `MED24`, `MED13L`), joined by the elongation kinase `CDK9`. Alongside it, the histone-modifying coactivators are lost: the acetyltransferase `CREBBP` and the shared core of the ASCOM/MLL3–MLL4 H3K4-methyltransferase complex^27^ — the nuclear-receptor coactivator `NCOA6` and `ASH2L` (the third core subunit `RBBP5` is retained by both baits). Chromatin remodellers (`ACTL6A`, `SS18L1`, the BAF subunit `SMARCC2`) and histone components (`H2BU1`, `H3-5`) complete the signature. Over-representation analysis echoes it (transcription-coactivator activity, the CORUM *Mediator complex, ASCOM* (MLL3/MLL4) and *BRD4* complexes; Supplementary Figure 2A), but we read it from the per-protein complex membership rather than the enrichment *p*-values, because affinity proteomics over-samples these abundant complexes — against the evaluated-proteome background only the *ASCOM* complex remains significant (Methods). Biologically, the coordinated loss of these ASCOM H3K4-methyltransferase contacts provides a direct, interaction-level explanation for the reduced promoter H3K4me3 levels documented in HD brain^14,15^, and complements the classical model of CBP/Mediator coactivator disruption^12,13^. These transcription-activation modules are resolved chiefly in the multi-epitope (Justice *et al*.) and human control-neuron (Gutiérrez-García *et al*.) datasets (Figure 2).

**Figure 3.**
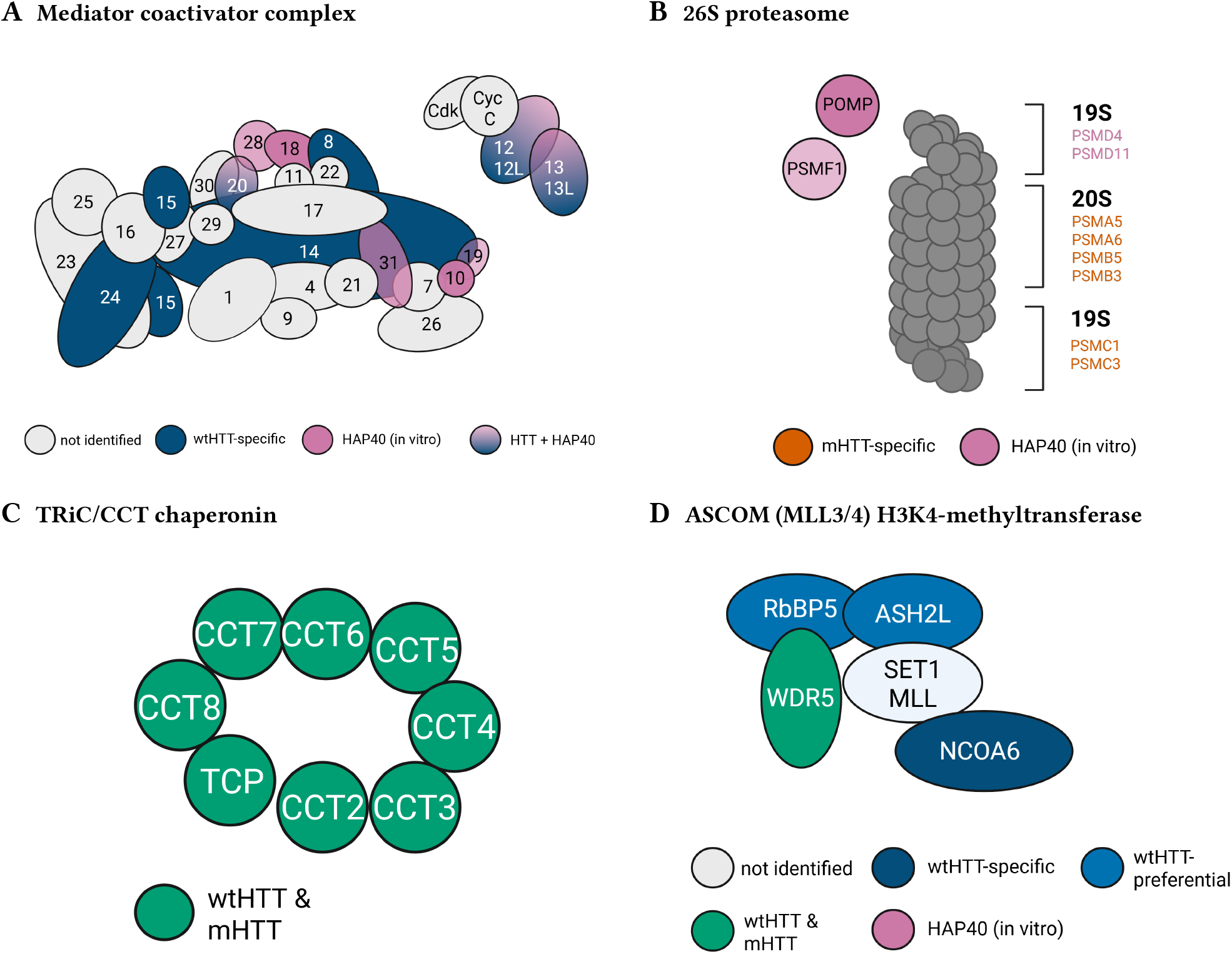
Signature complexes remodelled in (and conserved across) the HTT interactome, mapped onto their subunit topology. **(A)** The Mediator coactivator complex (subunits numbered MED#; kinase module — CDK8, cyclin C, MED12/12L, MED13/13L — at upper right). Six tail/head subunits are wtHTT-specific (dark blue; lost on expansion — `MED8`, `MED14`, `MED15`, `MED20`, `MED24`, `MED13L`), giving the interaction-level counterpart of the HTT-supported Mediator assembly weakened by the expansion. `MED15` is a single subunit drawn at its two structural contact points — its extended C-terminus forms two antiparallel helices wrapping around `MED14`^29^ — not two copies. **(B)** The 26S proteasome (20S proteolytic core capped by the 19S regulatory particle). mHTT-gained subunits (vermillion) span the 20S core (`PSMA5`, `PSMA6`, `PSMB3`, `PSMB5`) and the 19S AAA-ATPase ring (`PSMC3`) — the signature of a misfolding-prone bait recruiting the degradation machinery. **(C)** The cytosolic chaperonin TRiC/CCT (all eight subunits `CCT2`– `CCT8` and `TCP1`), engaged equally by wtHTT and mHTT (green, constitutive core) — the polyQ-independent interactor that serves as the internal positive control. **(D)** The ASCOM/MLL3–MLL4 H3K4-methyltransferase complex (the WRAD core — `WDR5`, `RBBP5`, `ASH2L` — with the SET1/MLL catalytic subunit and the nuclear-receptor coactivator `NCOA6`). `NCOA6` is wtHTT-specific (dark blue) and `ASH2L` wtHTT-preferential (blue), with `RBBP5` and `WDR5` in the constitutive core (green) — the H3K4-methyltransferase contacts lost on expansion, mirroring the Mediator pattern and providing an interaction-level basis for the reduced promoter H3K4me3 in HD^14^. The Mediator and proteasome panels additionally overlay contacts from the companion HAP40 interactome^10^ (mauve, bound by recombinant apo-HAP40 *in vitro* purple, contacted by both HTT and HAP40): in Mediator, `MED20` and `MED31` in the proteasome, the regulatory/assembly factors `PSMD4`, `PSMD11`, the maturation chaperone `POMP` and the inhibitor `PSMF1` (PI31), drawn as separate regulators rather than core subunits. Grey = not resolved in the HTT differential. Mediator topology after Justice *et al*.^23^; schematics created with BioRender.

**Figure 4.**
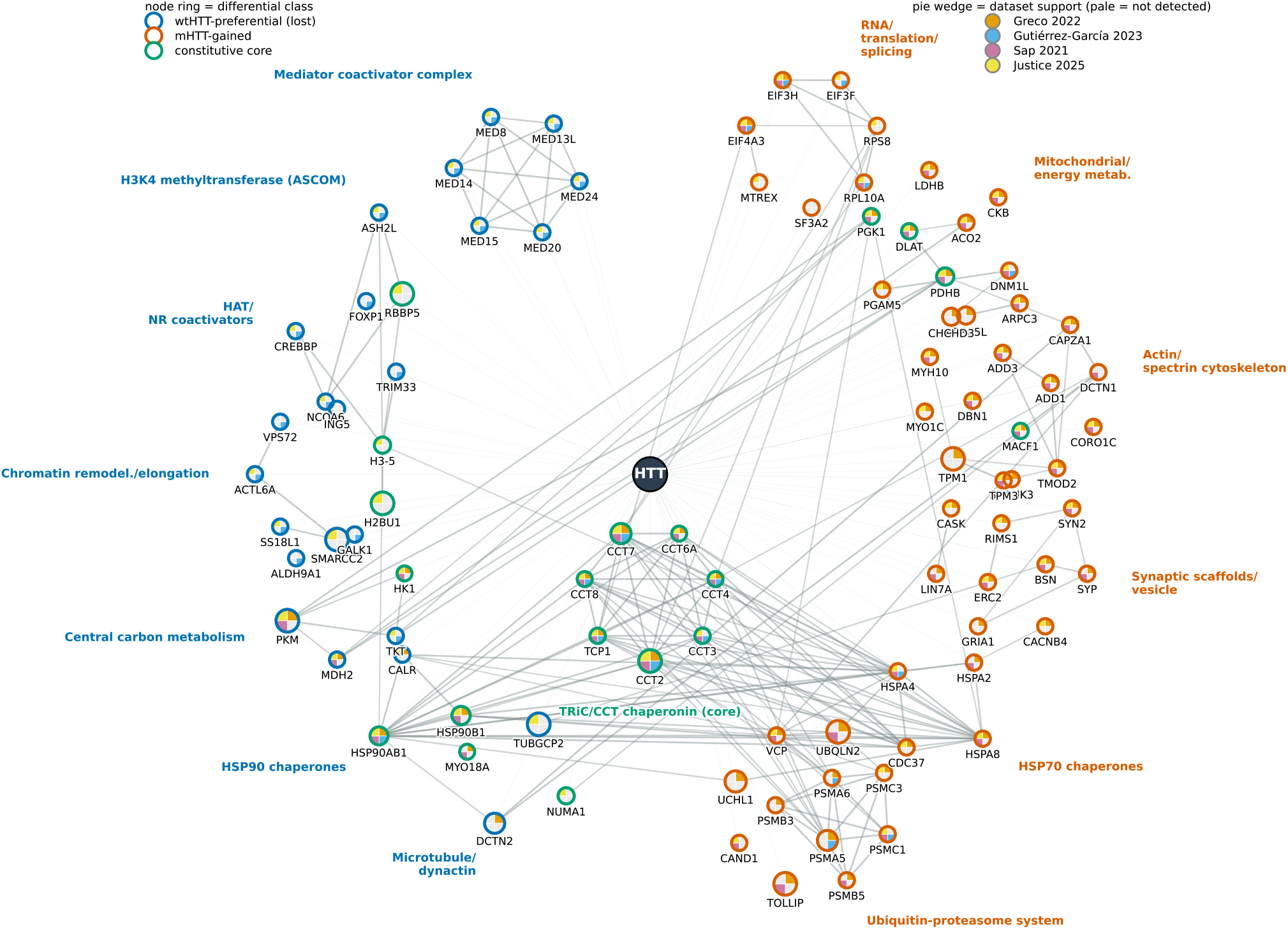
Dataset-resolved map of the remodelled HTT interactome. Curated functional-module members that received a condition-dependent or constitutive-core call, arranged around the HTT bait. Modules are grouped spatially — wtHTT-lost left, mHTT-gained right, the constitutive TRiC/CCT core at the centre. Each node’s *ring colour* encodes its differential class (blue, wtHTT-preferential; vermillion, mHTT-gained; green, constitutive core); the *pie wedges* encode source-dataset support (Greco 2022, Gutiérrez-García 2023, Sap 2021, Justice 2025; a wedge is filled in the dataset’s colour if that dataset detected the protein, pale grey otherwise); node size ∝ differential confidence (−log_10_ differential BFDR). Faint edges connect each interactor to the HTT bait. Colourblind-safe palette; legends inset.

A second wtHTT-preferential block comprised glycolytic and carbon-metabolism enzymes (`PKM`, `MDH2`, `DLD`, `TKT`, `HK1`, `GALK1` KEGG “carbon metabolism” and “pyruvate metabolism”), together with HSP90 chaperones (`HSP90AB1`, `HSP90B1`, `CALR`) and a microtubule/dynactin module (`DCTN2`, `ACTR1A`, `NUMA1`, `TUBGCP2`). The preferential association of energy-metabolism enzymes with wtHTT recovers a key observation of the Sap *et al*. dataset^24^ — itself one of the pooled inputs, so a consistency check rather than independent corroboration.

### mHTT gains proteostatic and synaptic contacts

The interactions gained by mHTT were both more numerous than the losses and, as individual calls, robustly supported: 182 of the 205 gains were recovered by the MAP point classifier (Supplementary Figure 10), so for this arm the decision-theoretic and point-estimate classifiers agree closely. The gained set was functionally distinct (Figure 1D; Supplementary Table 4). Its most robust complex-level signal was the *26S proteasome* — significant against both the genome and the evaluated-proteome background (CORUM, *p* ≈ 1 × 10^−4^ vs genome; Figure 3B) — and the mHTT-gained set further contained a broad ubiquitin–proteasome module (`PSMA5`, `PSMA6`, `PSMB3`, `PSMB5`, `PSMC3`, `UBQLN2`, `VCP`, `UCHL1`, `CAND1`) and HSP70 chaperones (`HSPA8`, `HSPA2`, `HSPA4`, `DNAJA2`, `CDC37`) — the expected signature of a misfolding-prone bait that recruits the degradation and disaggregation machinery^16,17^. Among the individual gains is the polyQdisease protein ataxin-2 (`ATXN2`)28, recovered specifically on the mHTT side. That these gains reflect condition-dependent contacts rather than non-specific background is supported directly: across studies the most *fragile* gains are the *least* promiscuously detected proteins — promiscuity here being detection breadth across *all* bait immunoprecipitations in the four source studies (a CRAPome-style measure of “sticky” background, computed over the many IPs of the four datasets rather than the two HTT baits alone; Supplementary Figure 6) — so per-call confidence and cross-study promiscuity are positively, not negatively, coupled (Supplementary Figure 7).

mHTT additionally gained an extensive synaptic and cytoskeletal interactome: presynaptic (`BSN`, `RIMS1`, `ERC2`, `SYP`, `SYN2`) and postsynaptic (`SHANK1`, `SHANK3`, `CASK`, `LIN7A`, `GRIA1`) scaffolds and a large actin/spectrin network (`ADD1`, `ADD3`, `MACF1`, `CAP1`, `ARPC3`, `TPM1`, `TPM3`, `VCL`). Against the genome these give the strongest GO terms overall (“synapse”, “postsynapse”, “glutamatergic synapse”; Supplementary Figure 2B), but this synaptic *over-representation* is detection-driven — it disappears against the evaluated-proteome background (only “actin filament binding” survives), so the synaptic signature should be read from the per-protein membership of these robustly-called contacts rather than from the genome-background enrichment *p*-values. A mitochondrial/TCA-cycle and translation set was also gained (`ACO2`, `DLAT`, `PDHB`, `DNM1L EIF3A`, `EIF3F`, `EIF3H`, `EIF4A3`). Notably, mHTT-gained engagement of translation factors including `EIF3H` again recovers a finding of the Sap *et al*. dataset^24^ (one of the pooled inputs), in which mutant huntingtin associated with the translation apparatus while wtHTT associated with energy metabolism — exactly the bidirectional pattern recovered here. These proteostasis, synaptic and cytoskeletal modules are carried most strongly by the mouse-striatal (Greco, Sap) and multi-epitope (Justice) datasets (Figure 2). Because soluble co-purification cannot by itself distinguish direct binding from indirect or co-aggregation–trapped association, these gained synaptic and cytoskeletal contacts are reported as condition-dependent engagement whose *directness* awaits orthogonal test.

### A constitutive core interactome is preserved on expansion

Complementing the remodelled sets, 157 proteins formed a robust constitutive core for which *both* the *maximum-a-posteriori* (MAP) classification *and* the decision-theoretic optimal call were *unchanged* — credible interactions with wtHTT and mHTT and no differential change (the 157 of the 294 unchanged calls whose point estimate also resolved to unchanged; Figure 1D). This constitutive core is the part of the HTT interactome that the polyQ expansion leaves intact, and it is dominated by HTT’s protein-folding and scaffolding machinery rather than its regulatory partners. The strongest complex was the cytosolic chaperonin TRiC/CCT — significant against both the genome and the evaluated-proteome background (CORUM “CCT complex”, *p* ≈ 4 × 10^−10^ vs genome; `CCT2`–`CCT8`, `TCP1` Figure 3C), recovered robustly in all four source datasets (Figure 2), and accompanied by “ATP-dependent protein folding chaperone” and “unfolded protein binding” enrichment (`HSP90AA1`, `HSPA5`, `HSPA9`, `HSPD1`, `DNAJA1`) and the Reactome terms “folding of actin by CCT/TriC” and “prefoldin-mediated transfer of substrate to CCT/TriC”. TRiC is the principal cellular chaperonin known to bind HTT and to suppress mHTT aggregation^30,31^ because it engages HTT largely through N-terminal determinants outside the polyQ tract, recovering it equally with both baits, as part of the constitutive core, is the expected behaviour and confirms that the analysis captures bona fide HTT interactions. The core further contained the obligate structural partner HAP40 (`F8A2`), the classic HTT palmitoyltransferase `ZDHHC17` (HIP14), the microtubule/ actin motor and scaffolding apparatus (`DYNC1H1`, `KIF5C`, `ARPC2`, `ARPC4`, `CAPZB`, `CDC42`, `RAC1`, `TLN1`), the translation/ribosomal machinery, and the R2TP/telomere-maintenance module (`RUVBL1`, `RUVBL2`, `WDR5`, `NPM1`). Thus the expansion spares HTT’s housekeeping engagement with its folding chaperonin, HAP40 and the transport apparatus, while selectively remodelling its transcription-regulatory (lost) and proteostatic/synaptic (gained) contacts. Notably, HAP40 (`F8A2`) — the obligate, deeply coevolved structural subunit that forms an exceptionally stable heterodimer with HTT^4,5^ — emerges from the same uniform analysis applied to every other protein, with no special handling, as one of the most robust core interactors, engaging wild-type and mutant HTT alike. The polyQ expansion therefore reshapes a *variable* interactome around this *constitutive* HTT–HAP40 scaffold; the HAP40 interactome itself and its stoichiometric signalling biology are addressed in a companion study^10^.

### HTT and HAP40 have largely distinct interactomes

Because a companion study^10^ profiled HAP40 directly — Strep-tagged in cells (*in vivo*) and as recombinant GST-HAP40 (*in vitro*) — and we scored those baits through the identical pipeline, we could ask how HTT’s interactome relates to that of its obligate partner^5,8^. The two are largely distinct: of the wtHTT and mHTT high-confidence partners (128 and 202 at *P* > 0.95), only 1 and 9 are shared with the *in vivo* HAP40-Strep interactome (232) — no more than expected by chance for sets of this size — and a substantial part of the HAP40 signal is **mitochondrial**. This separation is expected rather than surprising: only 12% of the over-expressed Strep-HAP40 is in complex with HTT^10^, so the *in vivo* HAP40 profile predominantly reports *apo*-HAP40 and largely reflects the contacts HAP40 makes outside the HTT heterodimer, dissected in the companion study^10^, and this separation is preserved across posterior cut-offs (Supplementary Figure 3). Consequently, the polyQ-driven remodelling of HTT’s contacts is — to the resolution of these data — largely **HAP40-independent**: the lost and gained sets intersect HAP40′s partners only modestly, so the disease-relevant rewiring is concentrated on the HTT face of the shared scaffold. Where the two proteins *do* contact the same complex they engage *distinct* subunits (at *P* > 0.95 no subunit is credibly bound by both; per-subunit engagement in Supplementary Figure 4, complex topologies in Figure 3); for the Mediator complex29, the few subunits contacted by recombinant HAP40 lie adjacent to those contacted by HTT (Figure 3).

### Coherence with the most comprehensive endogenous-HTT interactome

The richest single endogenous-HTT interactome in our pool — multi-epitope immunocapture of wild-type (Q20) and expanded (Q140) HTT across striatum, cortex and cerebellum23 — is also the most informative internal coherence check, for reasons independent of the result: it is the most comprehensive of the four datasets, the only input carrying orthogonal validation (thermal-proteome profiling, subcellular fractionation and *Drosophila* genetic-modifier assays), and it advocated the integrative direction this re-analysis takes. Because it also contributes a large share of the mutant-condition evidence (Supplementary Figure 11), agreement on the gain side reflects internal coherence rather than independent replication; the wtHTT arm and the divergences are accordingly the more informative comparisons. Read in that light, our single differential recovers several of its central observations. First, it mapped HTT onto nearly the entire Mediator complex (20/26 subunits) with preferential enrichment of the polyQ-rich tail subunit MED15, and showed by thermal-proteome profiling — orthogonal to co-purification, and so genuinely independent of our pooled evidence — that HTT stabilises Mediator and that mHTT redistributes MED15/MED27. That study also reports that mHTT continues to engage Mediator in an epitope- and tissue-dependent manner; our calling of six Mediator subunits (including `MED15`) as wtHTT-specific (Supplementary Table 4) therefore does not assert that mHTT never contacts Mediator, but that — unlike the wtHTT association — no credible mHTT–Mediator signal survives once the four datasets are combined, marking Mediator as an HTT-supported assembly the expansion destabilises rather than abolishes. Second, it captured the intact MLL3/MLL4 (ASCOM) H3K4-methyltransferase complex (`KMT2C`, `KDM6A`, `PAGR1`, `PAXIP1`, `NCOA6`) and reported its loss with mHTT; we independently call its shared core — `NCOA6` and `ASH2L` — wtHTT-preferential (with `RBBP5` retained by both baits). Third, on the gain side it found increased mHTT (Q140) interaction with acetyl-CoA/pyruvate-dehydrogenase proteins (`DLAT`, `PDHA1`, `PDHB`, `PDHX`) — we call `DLAT` and `PDHB` mHTT-gained — and its proteostasis (HSP70/chaperonin) and striatal genetic-modifier signatures are echoed by our mHTT-gained HSP70 module and `CACNB4`, and by the BAF/SWI-SNF members on the wtHTT side; these gain-side agreements are the ones partly anticipated by the shared samples.

The two divergences are more instructive than the agreements. CREBBP, reported there as detected largely independently of polyQ length, is called wtHTT-specific in our aggregate; and chaperone directionality is mixed (HSP70 mHTT-gained but HSP90 wtHTT-preferential). Both are expected consequences of averaging over tissue and epitope and of the dependence of condition-specific calls on detection sensitivity in the weaker bait, and mark interactions for which tissue-resolved follow-up is most warranted. The CREBBP call additionally bears on the classical CBP-sequestration model: because expanded polyQ is thought to sequester CBP through polyQ– polyQ contacts^12,13^, that model predicts a mHTT-*gained* interaction, whereas we recover CREBBP as wtHTT-specific. The two are reconcilable if sequestration occurs predominantly in the insoluble, aggregated phase that soluble co-purification does not sample: the expansion would then both *lose* the physiological soluble wtHTT– CBP association and *gain* an aggregate-phase one, two routes converging on the same depletion of available coactivator — a hypothesis our soluble data cannot test and that would require insoluble-fraction measurements to resolve.

## Discussion

Placing wtHTT and mHTT evidence inside one Bayesian model and asking, per protein, “in which bait is this interaction credible?” resolves the HTT interactome into two biologically distinct components. The expansion’s losses are dominated by HTT’s physiological coupling to the gene-activation apparatus — the Mediator/ARC coactivator complex, the ASCOM (MLL3/MLL4) H3K4-methyltransferase module, CREBBP, BRD4 and CDK9 — and by glycolytic/carbon-metabolism enzymes. Its gains are dominated by the ubiquitin–proteasome system and HSP70 chaperones, the polyQ protein ataxin-2, and a broad synaptic and actin-cytoskeletal network.

A methodological choice underpins these conclusions. Most published HTT interactomes are scored one study at a time, by fold-change thresholds or heuristic spectral-count scores (e.g. SAINT, CompPASS) that treat enrichment, reproducibility and specificity separately and cannot place wtHTT and mHTT on a common probability scale. BayesInteractomics instead integrates detection, enrichment and bait dose–response as explicit, dependency-aware probabilistic evidence, pools the heterogeneous protocols within one hierarchical model, and returns a posterior probability and a decision-theoretic call for every protein. Two advantages follow that are essential here: the wtHTT-versus-mHTT comparison contrasts two posteriors expressed on the same probability scale rather than incommensurable per-study statistics, and the explicit decision risk distinguishes a genuinely confident call from one favoured only at the point estimate. This pooling is consequential, not cosmetic: because the source studies overlap so little, 79% of the high-confidence (*P* > 0.95) interactors are supported by at most one study individually, and a conventional “reproducible in ≥2 studies” consensus would recover only 21% of them (Supplementary Figure 9) — yet where two or more studies do co-detect a protein, their effect magnitudes are statistically homogeneous (median *I*^2^ = 0, no significant heterogeneity in ∼ 95% of calls; Supplementary Table 3).

This architecture unifies observations usually treated in isolation. The coordinated loss of Mediator, CREBBP *and* the ASCOM H3K4-methyltransferase indicates that the expansion does not merely perturb one coactivator but uncouples HTT from the active-transcription machinery as a whole, furnishing an interaction-level basis for both the coactivator-disruption^12,13^ and the H3K4me3-loss^14,15^ phenotypes of HD. In parallel, the gained proteasome, UBQLN2/VCP and chaperone contacts are the expected behaviour of a misfolding-prone protein that recruits the proteostasis network^16–18^. This bidirectional pattern is not an artefact of our labelling: it recovers a within-study result of one of the pooled datasets24 (Figure 2) — energy-metabolism enzymes associating with wild-type HTT, and translation factors such as EIF3H with mutant HTT. Because that study is itself one of the four pooled here, this is a coherence check rather than independent replication.

Both arms are statistically robust, and the larger, better-supported arm is the set of gains. At the point estimate the MAP classification assigns the mHTT-gained direction to 351 proteins versus 46 in the wtHTT-lost direction, and 174 of the 197 mHTT-preferential calls are MAP-robust at a median differential false-discovery rate of effectively zero (Supplementary Figure 10). The gains are therefore not a low-confidence by-product of the decision rule but a genuine, condition-dependent expansion of HTT’s contacts that is as well resolved as — indeed slightly better resolved than — the coactivator losses. We tested directly whether the gains might nonetheless reflect non-specific, aggregation-associated detection rather than condition-dependent engagement, and found the opposite of what that model predicts: the most *fragile* gains are the *least* promiscuously detected proteins, and per-call confidence and cross-study promiscuity are positively — not negatively — coupled (Supplementary Figure 7). A non-specific “sticky background” would produce the reverse. The per-study effect magnitudes are also homogeneous rather than contradictory. The gains thus behave as reproducible, condition-dependent contacts; whether each reflects a direct or an indirect (for example co-aggregation–trapped) association is a distinct question that these soluble co-purification data cannot settle and that we carry as an explicit caveat. This is consistent with direct biophysical evidence that the polyQ expansion does not, of itself, change huntingtin’s affinity for its partners^32^ and that the expansion does not drastically alter HTT’s structure or properties^5^: the remodelling reported here is a redistribution of which contacts are engaged, not a wholesale change in binding chemistry.

The value of pooling is also evident in the structure of the evidence. The four source datasets overlap only modestly differences in species, tag, epitope, polyQ length and quantification mean each recovers a partial, noisy slice of the interactome (mean pairwise Jaccard 0.19; Supplementary Figure 6)^22–24,26^ — so a contact supported weakly but consistently across them registers as credible only once they are pooled in one combined probabilistic model. The residual per-protein scatter reflects under-powering, not disagreement: on the difference-in-differences contrast the model scores, the per-study effects are homogeneous (Cochran’s *Q*/*I*^2^: median *I*^2^ = 0, no heterogeneity for ∼ 95% of calls and all of the losses; Supplementary Table 3), yet no single study resolves a per-protein magnitude on its own, so only the variance-weighted model recovers it. We therefore report the *direction* of each differential call, not its per-protein effect size. Equally, this low overlap cautions that absence from any single study is weak evidence of a true non-interaction. More broadly, the integrated map carries a practical lesson: much work has pursued individual mHTT-*preferential* interactions as disease mechanisms or drug targets^3,12,13,16^ one study at a time, where aggregation artefacts are hard to separate from genuine engagement. Pooling lets the analysis confirm which contacts reproduce — most, here — and flag those that do not. `SH3GL3` (endophilin A3) is the instructive precedent: an early, influential mHTT-*preferential* interaction later shown to be polyQ-length-independent and aggregation-driven32, it is absent from our map (though, undetected in any pooled dataset, an absence of evidence rather than a positive exclusion). The map is thus as valuable for filtering out putative interactions that do *not* reproduce as for the contacts it nominates.

A further dimension concerns HAP40. The disease-relevant remodelling is concentrated on the HTT face of the shared scaffold, yet HTT and HAP40 repeatedly contact the *same* complexes (Mediator, the proteasome, the chaperonins) at distinct, spatially adjacent subunits — the pattern expected if the intact HTT–HAP40 heterodimer, not either subunit alone, is the physiological binding unit^10^. This carries a disease implication the cross-sectional differential cannot capture: because HAP40 and HTT abundances both decline in HD^10,11^ — the proposed stoichiometric HTT–HAP40 “rheostat” — the heterodimeric scaffold is itself depleted, so contacts scored here as HAP40-independent may nonetheless be lost through its collapse. The HAP40 interactome and its signalling biology are dissected in the companion study^10^; for the present map it suffices that HAP40′s abundance, not HTT’s alone, may shape the disease interactome.

Several limitations temper these conclusions. As a meta-analysis over orthologue-harmonised gene symbols, the map inherits the cell-type, tag and polyQ-length heterogeneity of its inputs and cannot, by itself, separate direct binding from indirect or co-aggregation–trapped association — a particularly important caveat for the mHTT-gained synaptic and cytoskeletal sets, where sequestration into inclusions can mimic interaction^32,33^. Although these gains are statistically robust and condition-dependent calls, robustness speaks to reproducibility of the differential signal, not to the directness or mechanism of the underlying contact. Indeed, all of our differential classes — and most strongly the constitutive core — are over-represented in the mHTT inclusion proteome^33^ (Supplementary Figure 8), although inclusion membership alone does not distinguish the gains from the losses. The condition-specific calls additionally depend on detection sensitivity in the weaker bait. Only the three mouse knock-in datasets^22–24^ profile both genotypes; the single human dataset^26^ (wild-type iPSC-derived striatal neurons) anchors the wtHTT interactome but, lacking an mHTT condition, contributes no mutant evidence. The mutant arm of the differential — and hence every mHTT-gained call — therefore rests on mouse models, to the resolution of these data, and awaits confirmation in human material. Orthogonal validation — reciprocal co-immunoprecipitation, proximity labelling at defined polyQ lengths, and stoichiometry-resolved structural work on the HTT–Mediator/ASCOM and HTT–proteasome interfaces^4^ — will be required to convert these high-confidence statistical calls into mechanism; for the HTT– Mediator association, Justice *et al*. have already provided initial orthogonal support by thermal-proteome profiling and reciprocal immunoprecipitation^23^. Nonetheless, the loss-plus-gain map nominates concrete, prioritised interfaces and ties the transcriptional and proteotoxic arms of HD to a single quantitative interactome — one layered upon a constitutive HTT–HAP40 core that the expansion leaves intact.

## Acknowledgements

We thank Todd M. Greco for kindly providing data, and Stefan Kochanek and Fabrice A. Klein for their support and helpful discussions.

## Methods

### Datasets and integration

Four published huntingtin affinity-proteomics datasets were integrated — Greco *et al*. 2022^22^, Justice *et al*. 2025^23^, Sap *et al*. 2021^24^ and Gutiérrez-García *et al*. 2023^26^ — together with the GST-HAP40 and HAP40-Strep comparator pull-downs. Protein identifiers were mapped to upper-case orthologous gene symbols, intensities were log_2_-transformed, and datasets were combined by full outer join on gene symbol. Because the protocols differ in dynamic range, abundances were normalised across protocols (median-of-ratios size factors plus per-protein cross-protocol row-centering, applied automatically when a protocol-scale mismatch was detected) before imputation. Missing values were then handled with a dropout-aware, missing-not-at-random (MNAR) imputation in which a per-column logistic dropout curve (detection probability as a function of intensity) drives a tilted-Gaussian sampler, so that proteins absent from a low-abundance arm are imputed consistently with the observed detection–intensity relationship rather than at a global mean. In plain terms: a protein that is missing simply because it is too scarce to detect in a low-abundance sample is filled in with a plausibly *low* value rather than an average one, so that a genuine non-detection is not turned into an artefactual mid-range signal.

### Bayesian interaction scoring

Each bait condition (wtHTT, mHTT, and the GST-HAP40 and HAP40-Strep comparators) was scored against the HTT bait (ENSP00000347184) with BayesInteractomics v1.2.0 (Julia 1.12.5); per condition, 4,338 proteins were evaluated from 49 control and 51–52 sample experiments. For every protein the framework integrates three complementary lines of evidence: a Beta–Bernoulli model of *detection* consistency (presence in bait versus control replicates), a hierarchical Bayesian model of *enrichment* (log_2_ fold-change, fitted by variational inference with parameters shared across the contributing protocols), and a robust Student-*t* linear-regression model of *dose–response* between prey and bait abundance (JZS Cauchy slope prior^34^, degrees of freedom selected by WAIC^35^). Rather than multiplying these as if independent — which inflates the combined evidence, because enrichment and detection are positively correlated under both hypotheses — their Bayes factors are combined by Bayesian model averaging over two dependency-aware models: a two-component (H0/H1) copula mixture with copula-family selection, and a three-component latent-class mixture fitted by expectation–maximization; the two are weighted by leave-one-out stacking36 with a 5% weight floor. Priors on the latent-class structure used an empirical-Bayes Dirichlet estimate (Minka fixed-point) marginalised over a BIC-weighted simplex grid. The combined posteriors were calibrated against parametric-simulation ground truth (an interaction-prevalence × effect-scale grid) by Platt scaling37 under an expected-calibration-error gate, yielding for each protein a posterior probability of interaction, a posterior error probability (PEP) and a local false-discovery rate; five automated input-quality checks (scale, replicate correlation, missingness asymmetry, intensity-distribution shape and PCA separation) were inspected beforehand. The model architecture, the deep-learning structural prior, the calibration against simulation ground truth and the head-to-head benchmarking against established AP-MS interaction scorers (median AUROC 0.747 on synthetic data with known truth) are developed and validated in the companion methods paper^25^. Credible interactors were defined at a Storey monotone step-down Bayesian false-discovery rate (BFDR) ≤ 0.0538 (196 for wtHTT, 297 for mHTT), consistent with hierarchical-mixture FDR control^39,40^. Throughout, “credible interactor” denotes this BFDR ≤ 0.05 set; a stricter per-protein posterior cut (*P* > 0.95; 128 for wtHTT, 202 for mHTT) is used only where set membership is compared directly — the HTT–HAP40 overlap analysis and the cross-study support tally (Supplementary Figure 9) — with results stable across the cut-off (Supplementary Figure 3).

### Differential analysis

The wtHTT-versus-mHTT contrast reports, per protein, the posterior probability of interaction in each condition (P_wtHTT_, P_mHTT_) and their posterior error probabilities, a differential Bayes factor and a standardised differential z-score, a *maximum-a-posteriori* (MAP) classification, and a decision-theoretic *optimal call* obtained by minimising the expected posterior decision risk over the actions {gained, reduced, condition-A-specific, condition-B-specific, unchanged, both-negative}. The expected risk of action *a* is ∑_*t*_ *P* (*t*) *L*(*a, t*) over the four truth states *t*, with *P* (*t*) the per-protein posterior over {gained, reduced, unchanged, both-negative} (renormalised from the per-class posterior error probabilities) and *L* the default asymmetric loss matrix below (rows = action, columns = truth state, in that order; zero on the diagonal):

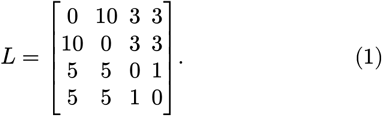

A direction reversal (calling *gained* when the truth is *reduced*, or vice versa) is penalised most heavily (10, as reversing a published direction is costlier than missing one); a missed condition-dependent call (*unchanged*/*both-negative* when the truth is *gained*/*reduced*) costs 5, an overclaim (a directional call when the truth is *unchanged*) costs 3, and a slip within the no-interaction region costs 1. The two condition-specific actions lie outside this 4×4 matrix and are assigned by coverage (a credible interactor of one bait only). We define interaction direction by the decision-theoretic optimal call: the “gained” calls (*n* = 50) and the condition-A-specific calls (*n* = 20) together form the *wtHTT-preferential* set lost on expansion (*n* = 70); the “reduced” calls (*n* = 197) and the condition-Bspecific calls (*n* = 8) form the *mHTT-gained* set (*n* = 205). The differential z-score (Δz = z_wtHTT_ − z_mHTT_, a raw difference in log-enrichment) and the posterior comparison P_wtHTT_ vs P_mHTT_ are reported alongside and are broadly concordant with the call, but the optimal call — not the sign of Δz or a strict posterior inequality — is the operative criterion, because a minority of proteins have a posterior saturated at ≈ 1 in both baits or credible in only one. The shared core (*n* = 157) comprises the proteins for which both the MAP classification and the optimal call were *unchanged* (the robust subset of the 294 unchanged calls; the bait HTT excluded). As a robustness tier, we intersected each directional decision-theoretic call with the MAP classification, retaining proteins for which both resolved to the same class (MAP-robust: 25/50 wtHTT-preferential, 20/20 wtHTT-specific, 174/197 mHTT-preferential, 8/8 mHTT-specific). The decision-theoretic optimal calls — the principled Bayes action under the full posterior — are the primary analysis; the MAP intersection is reported as a conservative confidence check. The mHTT arm of this contrast derives only from the three genotype-split datasets^22–24^ the human dataset^26^ profiles wild-type HTT only and so informs the wtHTT arm but adds no mHTT evidence.

### Robustness and cross-study support

The robustness, cross-study-support and sensitivity analyses that underpin the confidence and reproducibility claims of the Results and Discussion were all computed in Python with NumPy and SciPy^41^ from the committed differential table and the raw and MNAR-imputed matrices, and none refit the model. *Call robustness*. For each directional class we recomputed the fraction of members retained as the Bayesian-FDR acceptance threshold τ was tightened from 0.5 towards 0 under two distinct criteria — the per-condition interaction BFDR (a genuine interactor of the engaged bait) and the differential BFDR (condition-dependent rather than unchanged) — and cross-tabulated every call on two orthogonal axes, statistical robustness (MAP classification agrees with the optimal call) and cross-study breadth, to form two-track support tiers anchored by the constitutive core (Supplementary Figure 10). *Cross-study detection support*. Detection was scored per source study from the raw matrix as presence in ≥ 50% of that study’s bait immunoprecipitations. From this we computed, for every condition-dependent call, its detection breadth (number of supporting studies), the pairwise Jaccard index42 and exclusive intersections (UpSet) of the four studies’ detected proteomes, and a leave-one-study-out check — attributing each single-study call to its sole supporting study — confirming that no single dataset is load-bearing (Supplementary Figure 6). *Model-refit leave-one-study-out*. As a stronger test than this evidence-level check, the full pipeline — both single-condition models and the wtHTT-versus-mHTT differential — was re-fit four times with BayesInteractomics (v1.2.1; *Julia*, 16 threads), each time removing one source study’s protocol(s) from the pooled matrix, and each fold’s directional calls were compared protein-by-protein to the full-data calls (Supplementary Figure 11). *Value of integration*. To show that integration recovers signal no single study or reproducibility filter would, each credible interactor (*P* > 0.95) was re-scored by a conventional per-study rule (detected in ≥ 50% of a study’s bait IPs and ≥ 2-fold enriched over that study’s controls) and the fraction recovered by a “reproducible in ≥ 2 studies” consensus tallied; for the harder differential question a fair naive baseline computed a Welch mHTT-versus-wtHTT *t*-test within each genotype-split study on the same MNAR-imputed matrix, combined across studies by sample-size-weighted Stouffer’s *Z*43 (and by direction-consistent vote-counting) under Benjamini–Hochberg FDR control44 (Supplementary Figure 9). *Evidence composition and promiscuity*. For each condition-dependent protein the evidence for its engaged bait was decomposed into the three component log_10_ Bayes factors (detection, enrichment, dose– response), compared between the gained and lost classes by the Mann–Whitney *U* test45 and expressed as each arm’s mean fractional contribution to the positive log-evidence (Supplementary Figure 5). As a study-balanced promiscuity measure we used a CRAPome-style detection breadth46 — the fraction of a study’s IPs detecting the protein, averaged with equal weight across the four studies — with mean log_2_ abundance as a control. To test the “sticky background” objection that the least-confident gains might be non-specific contaminants, this promiscuity measure was stratified by support tier (fragile = the *neither* tier, neither MAP-robust nor detected in ≥2 studies) and compared between fragile and supported calls within each direction by the Mann–Whitney *U* test, with the per-call number of satisfied support axes correlated against promiscuity by Spearman’s ρ^47^ (Supplementary Figure 7). *Effect-size homogeneity*. To test whether the per-study effect magnitudes are inconsistent or merely underpowered, each call’s genotype effect was expressed on the difference-in-differences contrast the model scores — enrichment over the genotype-matched control in mHTT minus the same in wtHTT (reducing to a bait-only contrast for the single-pooled-control Sap dataset) — with a standard error propagated from the replicate variances, and tested for cross-study heterogeneity with Cochran’s *Q*^48^ and the *I*^2^ statistic^49^ across the studies detecting the protein (≥ 50% of bait IPs in ≥ 2 of the three genotype-split datasets), on both the MNAR-imputed and the raw observed-only matrices; the model’s variance-weighted magnitude (|Δ*z*|) was correlated with the studies’ mean |DiD| by Spearman’s ρ (Supplementary Table 3). *External validation and sensitivity*. Each differential class was tested for over-representation among proteins sequestered into mHTT inclusions^33^ by Fisher’s exact test^50^ against the evaluated-proteome background (one-sided), with gains compared to losses directly (twosided; Supplementary Figure 8). Finally, the four-bait interactor counts and their pairwise Jaccard overlaps were recomputed as the posterior cut-off τ was swept from 0.5 to 0.999, confirming the *P* > 0.95 comparisons are not special points (Supplementary Figure 3).

### Enrichment and visualization

Functional over-representation used g:Profiler (g:GOSt, *Homo sapiens*) against GO, KEGG, Reactome and CORUM with the g:SCS correction at *α* = 0.05^51^. Because affinity proteomics samples the proteome non-uniformly, we computed over-representation against two backgrounds: the default genomewide annotated set and a custom background of the 4,338 evaluated proteins (the detectable universe). The detectable universe is itself strongly over-represented, relative to the genome, for synaptic, proteasomal, chaperone, ribosomal and transcription-machinery terms (e.g. synapse *p* ≈ 10^−202^; 25 of 26 core-Mediator subunits detectable), so genome-background over-representation of the differential subsets largely reflects detectability rather than condition-specific selectivity. We therefore base the functional interpretation on curated complex membership and the per-protein interaction calls, present the genome-background enrichment as descriptive (Supplementary Figure 2), and use the evaluated-proteome background as the specificity test: against it the constitutive TRiC/CCT core and the gained 26S proteasome remain significant, whereas the broad synaptic and transcription-machinery terms do not. Complex- and pathway-level terms were prioritised for interpretation. All figures were rendered with the Lilaq library for Typst directly from the derived data.

### Data and code availability

This study re-analyses four previously published affinity-proteomics datasets; their raw data are available from the original publications and associated repositories^22–24,26^. All derived data generated here — the source differential-interactome table, the call-specific protein lists (also provided as the per-call sheets of *supplementary_tables*.*xlsx*), the derived enrichment results and all per-figure source data — are archived on Zenodo (DOI: 10.5281/zenodo.21219196), where a snapshot of the analysis and figure-generation code is deposited alongside them. The BayesInteractomics framework used for interaction scoring is openly developed at github.com/ma-seefelder/BayesInteractomics.jl — with the version used here archived in the same Zenodo record — and described in the companion methods paper^25^.

## Supplementary information

**Supplementary Table 1.** Source datasets integrated by the meta-analysis. The four published huntingtin affinity-proteomics studies pooled by BayesInteractomics, with their species and tissue of origin, HTT bait variants (polyQ length and age where applicable), capture chemistry and the number of bait and control immunoprecipitations (IPs), together with the two directly-profiled HAP40 comparator baits from the companion study^10^. Sample-size details are as reported in the source publications.

| Dataset | Species / tissue | HTT baits (polyQ; age) | Capture chemistry | IPs (bait / control) | Ref. |
| --- | --- | --- | --- | --- | --- |
| **Greco *et al.* 2022** | Mouse striatum | Q20, Q140 knock-in; 2 & 10 mo | FLAG immunoaffinity purification-MS (label-free + metabolic labelling) | 12 / 12 | <sup>22</sup> |
| **Justice *et al.* 2025** | Mouse striatum, cortex, cerebellum | Q20, Q140; 8 & 40 wk | Multi-epitope immunocapture (2B7/4C9 N-term, 3E10/4E10 central) | 81 / IgG | <sup>23</sup> |
| **Sap *et al.* 2021** | Mouse cortex | Control, Q20, Q140; 4 replicates each | Chemical-crosslinking FLAG co-immunoprecipitation | 8 / 4 | <sup>24</sup> |
| **Gutiérrez-García *et al.* 2023** | Human iPSC-derived striatal neurons | Endogenous total HTT (wild-type) | Label-free co-immunoprecipitation-MS | 3 / 3 (GFP) | <sup>26</sup> |
| **HAP40-Strep (companion)** | Cells (*in vivo*, complex-capable) | Strep-tagged HAP40 | Strep affinity capture | — | <sup>10</sup> |
| **GST-HAP40 (companion)** | Recombinant (*in vitro*, direct binders) | GST-HAP40 | GST pull-down | — | <sup>10</sup> |

**Supplementary Table 2.**
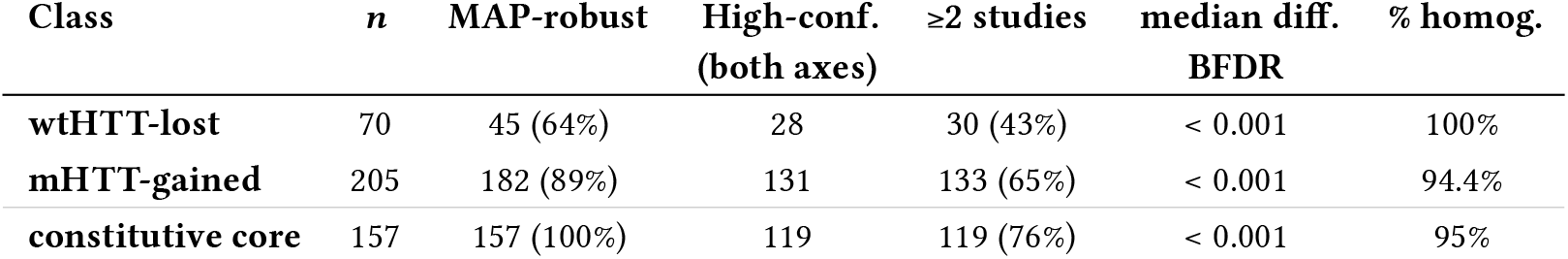
Both arms of the condition-dependent interactome are statistically well supported. Per-direction confidence summary for the 70 wtHTT-lost and 205 mHTT-gained directional calls, with the 157-protein constitutive core as the high-confidence anchor. *MAP-robust*, the decision-theoretic optimal call is also recovered by the maximum-a-posteriori point classification; *high-confidence (both axes)*, MAP-robust *and* detected in ≥2 of the four source studies (the gold tier of Supplementary Figure 10); *≥2 studies*, cross-study detection breadth (Supplementary Figure 6); *median diff. BFDR*, the median differential Bayesian false-discovery rate over the calls with a defined value (saturated/single-bait calls excluded); *% homog*., the fraction with no cross-study effect-size heterogeneity (Supplementary Table 3). The larger mHTT-gained arm is the more robust of the two — 182 of its 205 calls are MAP-robust and 131 reach the both-axes high-confidence tier, against 45 and 28 of the 70 losses — overturning the earlier impression that the gains were individually low-confidence.

| Class | *n* | MAP-robust | High-conf.<br>(both axes) | $\geq 2$ studies | median diff.<br>BFDR | % homog. |
| --- | --- | --- | --- | --- | --- | --- |
| wtHTT-lost | 70 | 45 (64%) | 28 | 30 (43%) | < 0.001 | 100% |
| mHTT-gained | 205 | 182 (89%) | 131 | 133 (65%) | < 0.001 | 94.4% |
| constitutive core | 157 | 157 (100%) | 119 | 119 (76%) | < 0.001 | 95% |

**Supplementary Table 3.**
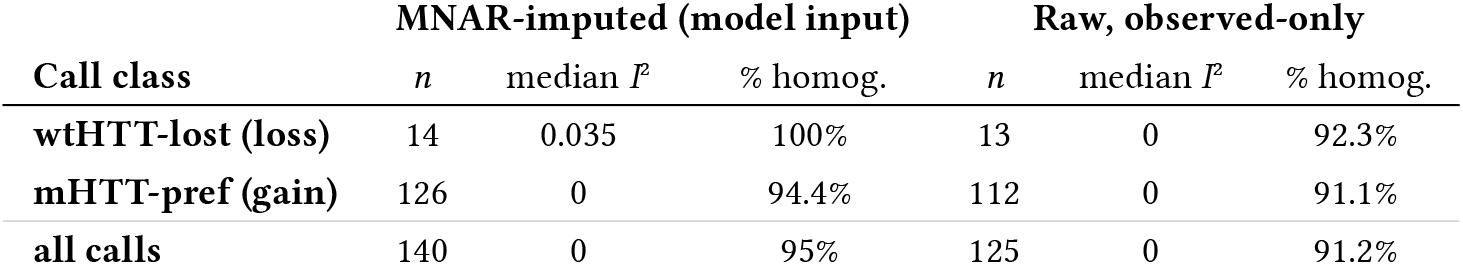
Per-study effect-size magnitudes are consistent across studies — underpowered, not contradictory. For every directional call co-detected (in ≥ half the baits) in at least two of the three genotype-split source studies, the genotype effect was expressed on the *difference-in-differences* contrast the model scores (enrichment over the genotype-matched control in mHTT minus the same in wtHTT) and tested for cross-study heterogeneity with Cochran’s *Q*^48^ / *I*^249^ on each study’s point estimate ± its standard error. Median *I*^2^ = 0 and ∼ 95% of calls (all of the wtHTT-losses) show no heterogeneity beyond sampling error (*Q p* ≥ 0.05): the studies do not disagree on magnitude — the ∼ 7-fold spread of the raw point estimates is sampling noise at the *n* = 4–6 replicates per study. The result is unchanged on raw, non-imputed intensities (right block), so it is not an artefact of the MNAR imputation. No single study resolves a per-protein magnitude on its own (only 1 call is individually significant in two studies at once); only the variance-weighted model recovers the aggregate effect — its |Δ*z*| tracks the studies’ mean |DiD| (Spearman^47^ ρ = 0.406, *p* ∼ 10^−9^). We therefore report the *direction* of each differential call, not its per-protein effect size. Sap 2021 has a single pooled control, so its DiD reduces to a bait-only contrast; *n* is the number of calls testable in each treatment.

**Supplementary Table 4.**
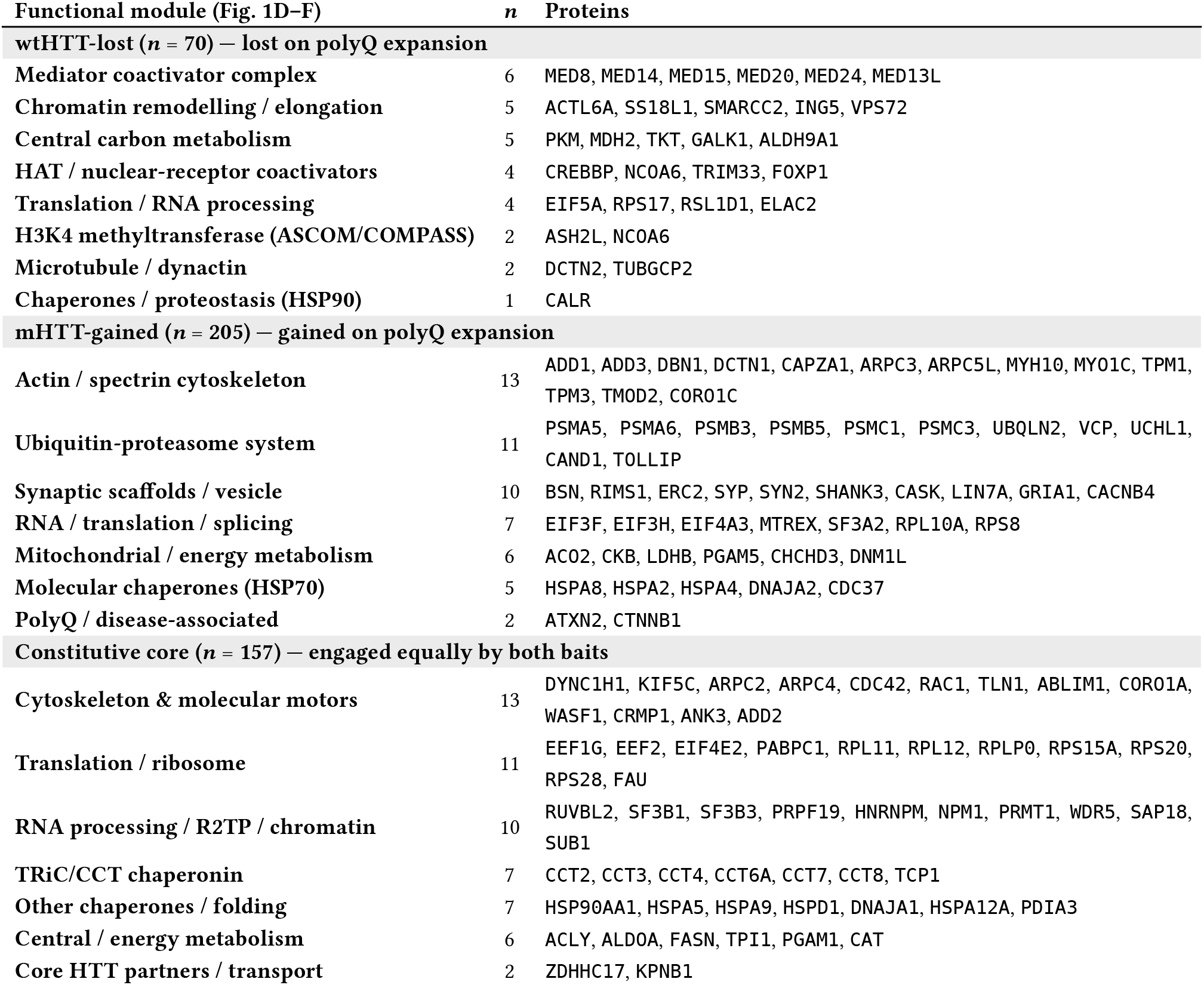
Functional-module composition of the differential HTT interactome, grouped by call. Curated functional-module membership (the categories of Figure 1 D–F) for the three calls — wtHTT-lost (lost on polyQ expansion), mHTT-gained (gained on polyQ expansion) and the constitutive core. *n* is the number of called proteins in each module; modules are ordered by size within each call. Only proteins assigned to a curated module are listed here; the *complete* per-protein call lists (all 70 / 205 / 157 proteins, with posterior probabilities, differential z-scores and Bayesian false-discovery rates) are provided as the per-call sheets of the accompanying *supplementary_tables*.*xlsx*, and the per-protein values are visualised in Supplementary Figure 1.

**Supplementary Figure 1.**
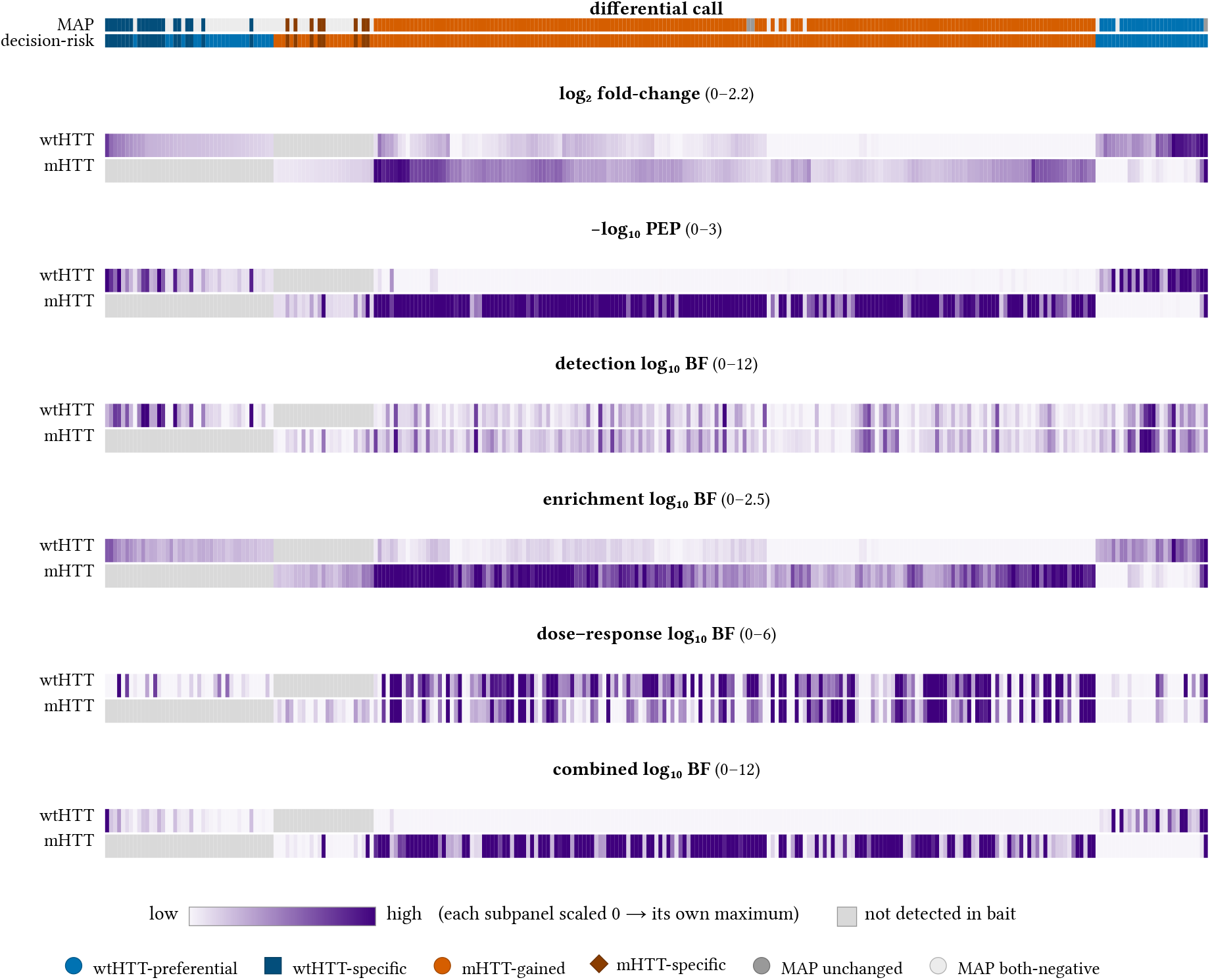
Per-metric map of the condition-dependent interactome, wtHTT versus mHTT. All 275 condition-dependent proteins (columns) are ordered once by hierarchical clustering (Ward linkage) of their two-dimensional log_2_ fold-change profile and held in that order across every subpanel; clustering separates a wtHTT-high block (lost on expansion) from an mHTT-high block (gained). The two *call* strips give the differential class under the point-estimate MAP classification and under the decision-risk-minimised optimal call — the two largely agree for both arms (most mHTT-gained proteins are MAP-concordant under the corrected differential). Each subsequent subpanel shows, for the *wtHTT* and *mHTT* baits, the per-protein log_2_ fold-change (bait vs control), the posterior error probability (as −log_10_ PEP) and the detection, enrichment, dose–response and combined log_10_ Bayes factors. Within every subpanel the colour scales from pale (no support) to saturated purple (strong support), normalised to that subpanel’s own maximum (given in its title); grey = not detected/not credible in that bait. The complete per-protein values for every protein are listed in Supplementary Table 4 and, per protein, the per-call sheets of *supplementary_tables*.*xlsx*.

**Supplementary Figure 2.**
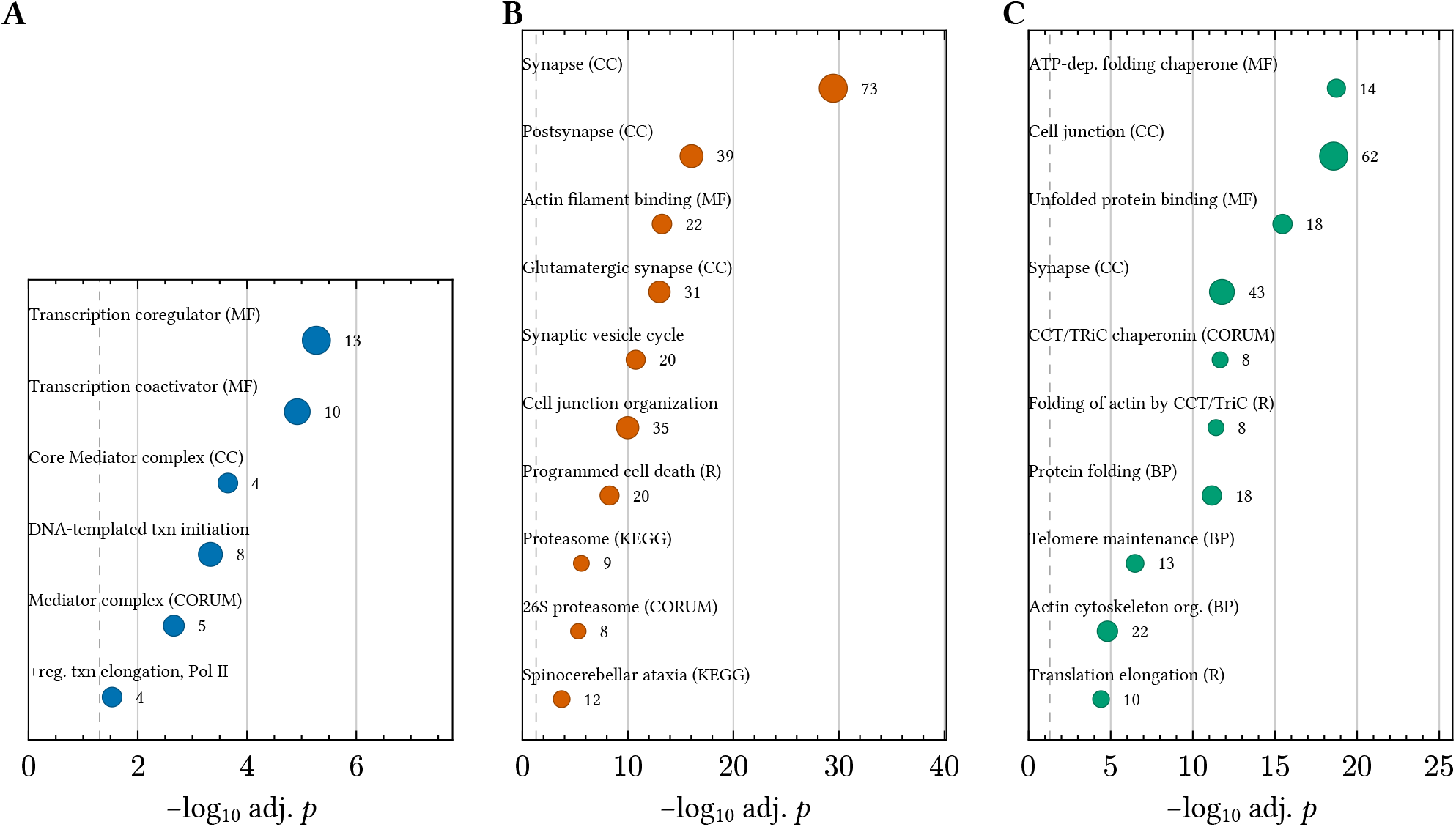
Functional over-representation of the differential interactome sets (genome background). g:Profiler^51^ over-representation analysis for **(A)** the wtHTT-lost set (*n* = 70), **(B)** the mHTT-gained set (*n* = 205) and **(C)** the constitutive core (*n* = 157). Each dot is a term: x-axis, −log_10_ g:SCS-adjusted *p*-value (dashed line, *p* = 0.05); dot area ∝ set proteins annotated (count beside each dot); GO, KEGG (K), Reactome (R) and CORUM sources indicated. wtHTT-lost contacts are dominated by the transcription-activation machinery and carbon metabolism; mHTT-gained contacts converge on the synapse, the actin cytoskeleton, the 26S proteasome and the polyQ “spinocerebellar ataxia” module; the core is dominated by the TRiC/CCT chaperonin and folding/ transport machinery. These panels use the whole-genome background and are descriptive: because the detectable proteome is itself enriched for these abundant categories, against the evaluated-proteome background only the ASCOM complex (A), the 26S proteasome and actin binding (B) and the TRiC/CCT core (C) remain significant (Methods). The functional claims in the main text therefore rest on the per-protein complex membership (Figure 1D) rather than on these *p*-values.

**Supplementary Figure 3.**
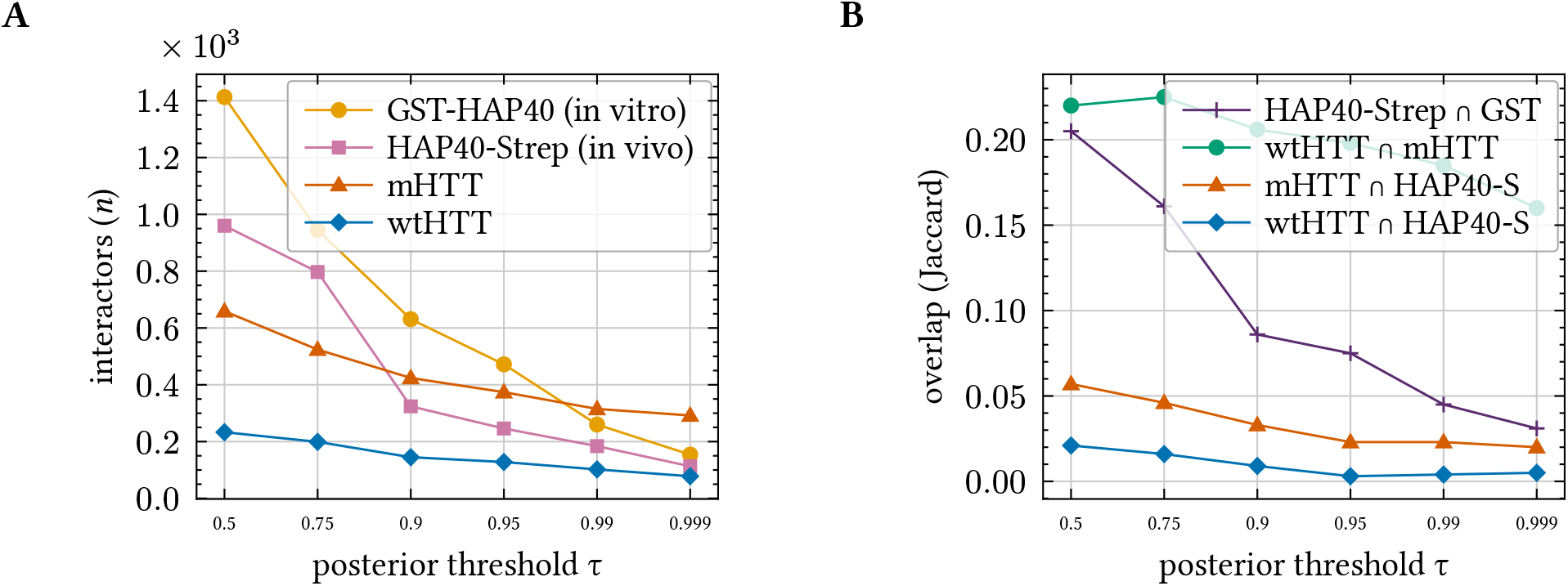
Robustness of the four-bait comparison to the posterior-probability cut-off. **(A)** Number of credible interactors per bait and **(B)** pairwise overlap (Jaccard index) as the posterior threshold τ is swept from 0.5 to 0.999. The relative ordering of the interactomes and the HTT/HAP40 overlap structure are preserved across the range, so the *P* > 0.95 cut-off used for the HTT/HAP40 comparison in the main text is not a special point. wtHTT and mHTT are kept separate throughout.

**Supplementary Figure 4.**
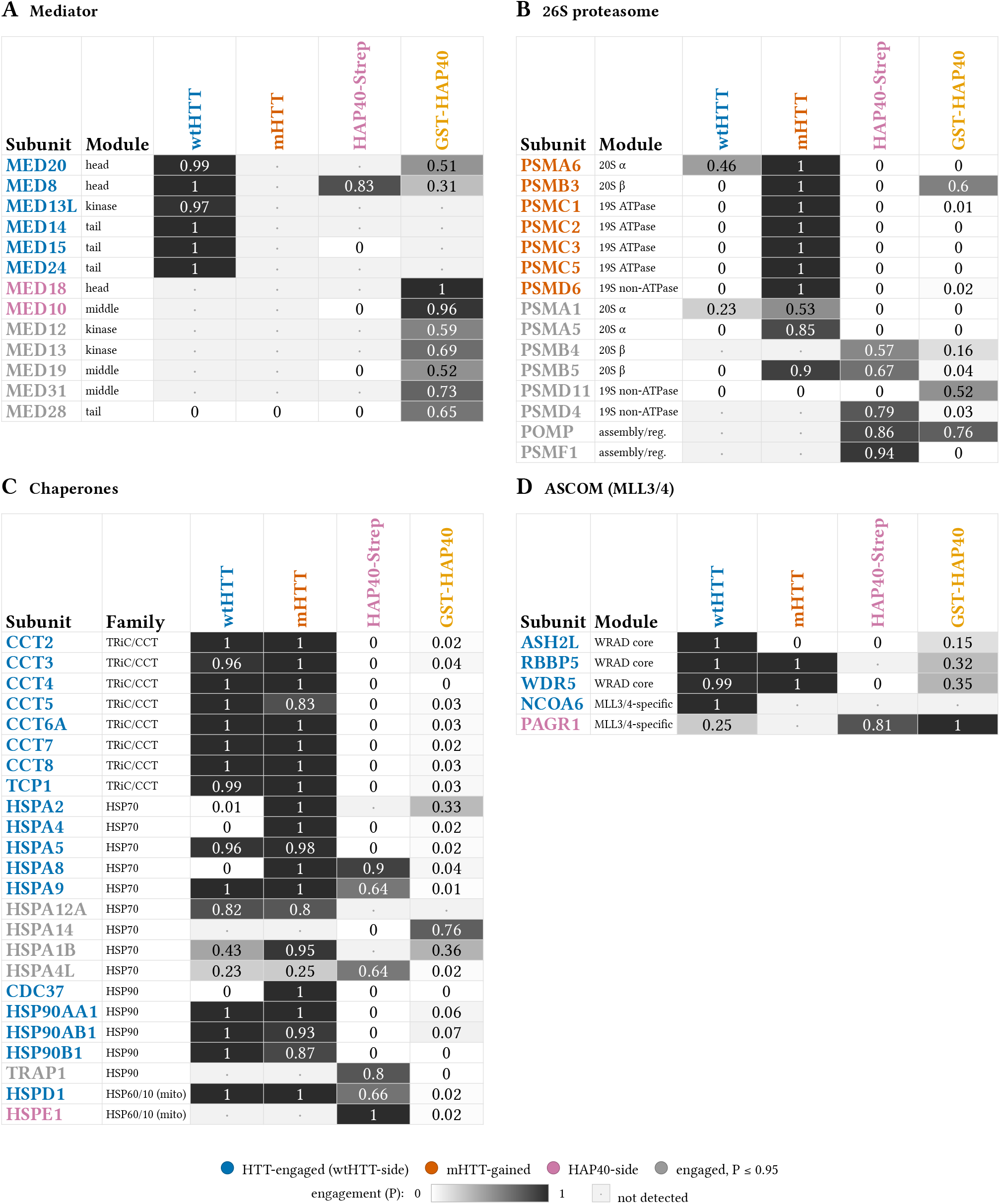
Complete per-subunit engagement of the four signature complexes across the four baits. Per-subunit basis for the HTT–HAP40 comparison in the Results (where HTT and HAP40 share a complex, they engage distinct subunits): **(A)** the Mediator coactivator complex, **(B)** the 26S proteasome, **(C)** the chaperone families (TRiC/CCT, HSP70, HSP90, mitochondrial HSP60/10) and **(D)** the ASCOM/MLL3–MLL4 H3K4-methyltransferase complex. Each cell, posterior probability of interaction for that subunit with that bait (greyscale key); subunit labels coloured by the side engaging them credibly at *P* > 0.95 (blue, HTT/wtHTT-side; vermillion, mHTT-gained; mauve, HAP40-side; grey, engaged but *P* ≤ 0.95 in every bait); “·” = not detected with that bait.

**Supplementary Figure 5.**
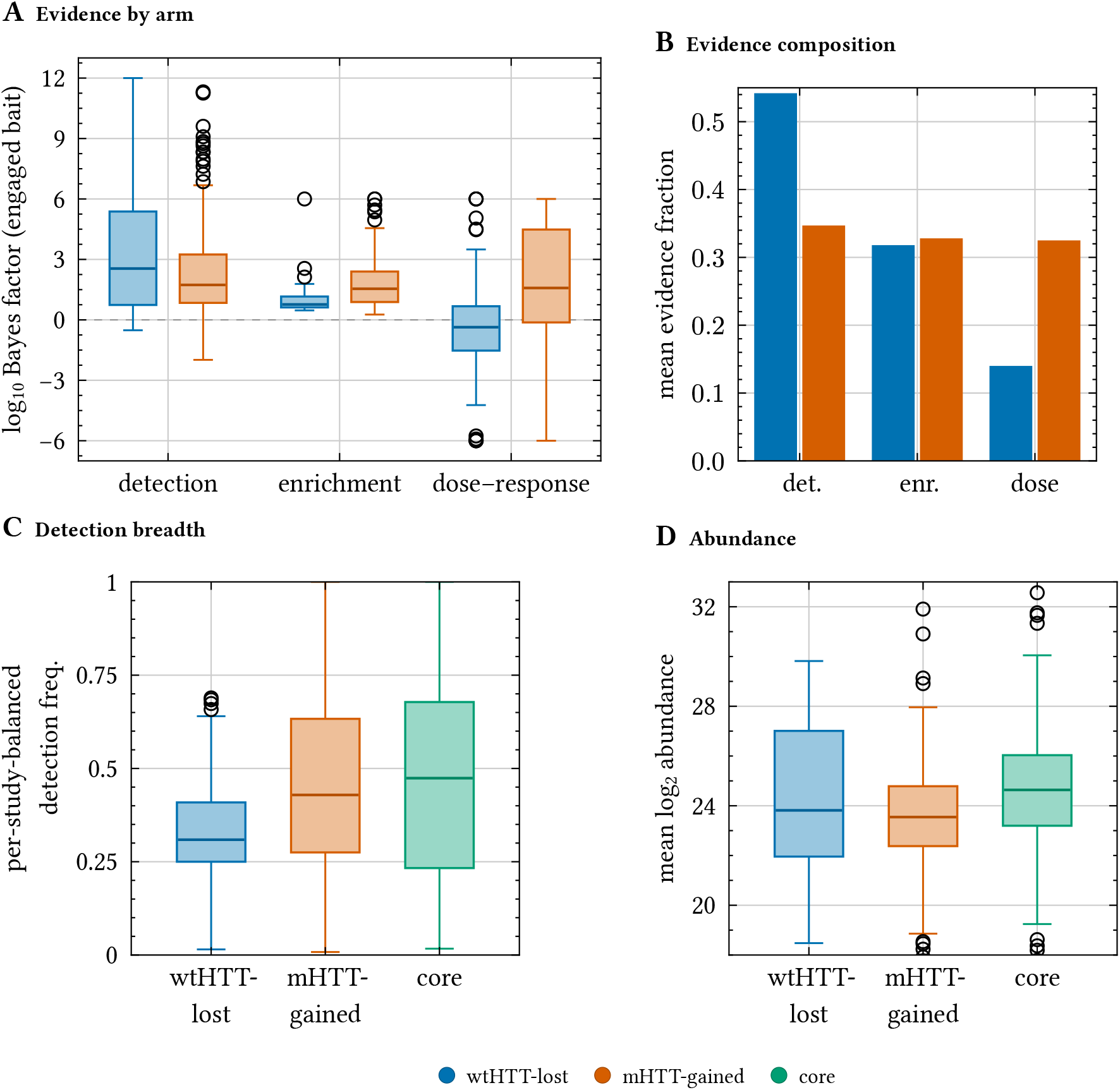
The mHTT-gained and wtHTT-lost interactomes draw on differently-composed evidence. **(A)** Distribution of the three per-protein evidence Bayes factors — detection, enrichment and dose–response (prey–bait abundance correlation) — for the bait each condition-dependent protein is called for (log_10_ scale; the correlation BF is model-capped at ±10^6^). All three arms differ between the classes (Mann–Whitney^45^ detection *p* = 0.0402, enrichment *p* < 10^−4^, dose–response *p* < 10^−4^): the wtHTT-losses carry the higher detection evidence (median 2.546 vs 1.734) while the mHTT-gains carry the higher enrichment (1.543 vs 0.764) and dose–response (1.582 vs −0.365) evidence. **(B)** Mean fractional contribution of each arm to a protein’s positive log-evidence: wtHTT-losses are detection-dominated (54% detection), whereas the mHTT-gains draw more evenly on detection, enrichment and dose–response (35/33/33%). **(C)** CRAPome-style^46^ detection breadth — the fraction of a study’s bait IPs detecting each protein, averaged with equal weight across the four studies (removing the dataset-size confound): mHTT-gains are somewhat more broadly detected than wtHTT-losses (median 0.429 vs 0.309, *p* = 0.0004), but this is a class-level difference; *within* the gains the least-confident calls are the least promiscuous (Supplementary Figure 7). **(D)** Mean log_2_ abundance: mHTT-gains are *not* more abundant (if anything marginally lower), so the breadth in (C) is not an abundance artefact. Boxes show the quartiles and median; whiskers, 1.5×IQR; points, outliers.

**Supplementary Figure 6.**
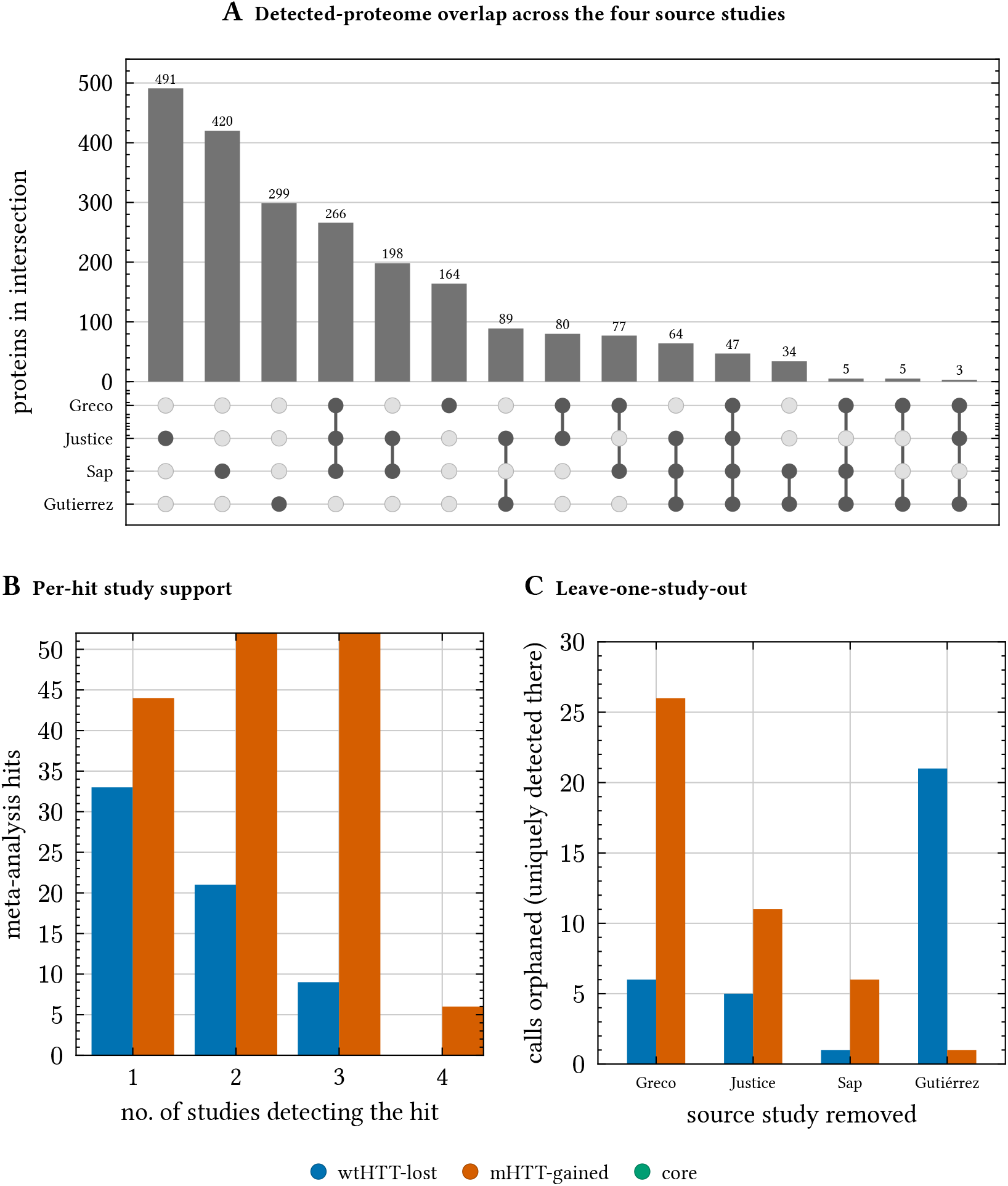
The source datasets overlap only modestly; the meta-analysis recovers interactors no single study would, yet no single study is load-bearing. **(A)** UpSet plot of the proteins detected (in ≥50% of a study’s bait immunoprecipitations) by each source study: bars give the size of each exclusive intersection, the dot-matrix its study membership. The three largest intersections are *single-study* (Justice 491, Sap 420, Gutiérrez-García 299 unique); only 47 proteins are detected by all four. Mean pairwise Jaccard index^42^ = 0.194. **(B)** For the condition-dependent hits, the number of studies that detect each: 48% of the wtHTT-lost set (33/69) is seen by only one study, whereas mHTT-gains are detected more broadly (peak at three studies; Supplementary Figure 5C). **(C)** The complementary leave-one-study-out check: attributing every single-study hit to the study that detects it, the most any one study *uniquely* contributes is 26 calls (Gutiérrez-García, whose endogenous-HTT capture detects a distinct subset),so removing any single study leaves the differential call set essentially intact. No single dataset reconstructs the interactome — yet none dominates it — motivating the integrated analysis.

**Supplementary Figure 7.**
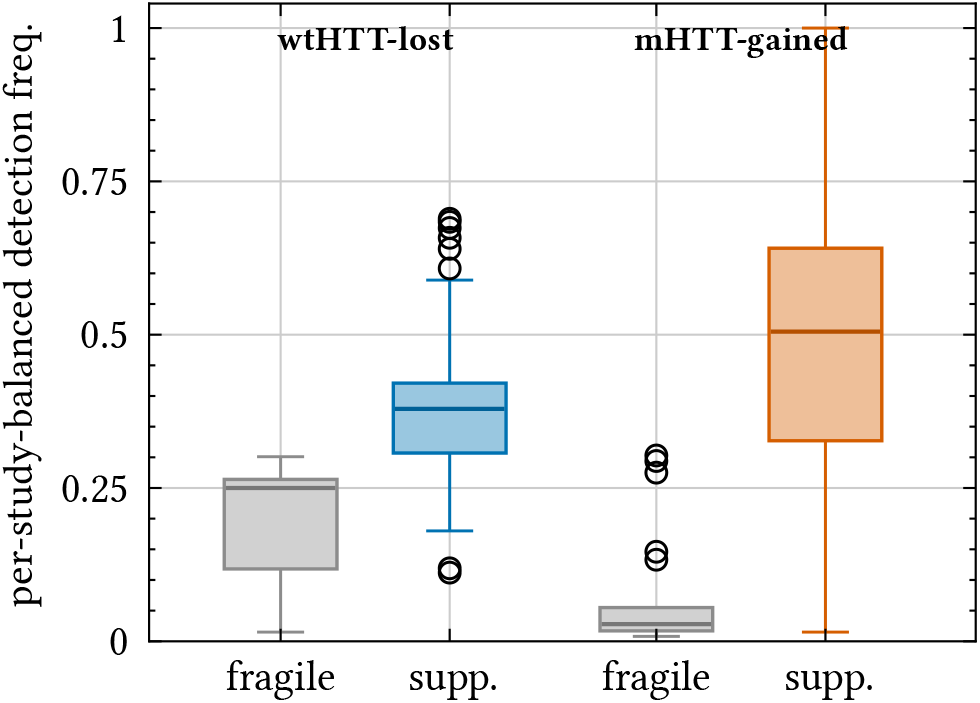
The least-confident gains are the *least* promiscuously detected — the inverse of a non-specific “sticky background”. A common objection to gain-of-interaction calls is that the weakest of them are non-specific contaminants — abundant, “sticky” proteins detected indiscriminately across pull-downs. If so, the lowest-confidence gains should be the *most* promiscuously detected. The data show the opposite. Each directional call is split into *fragile* (the *neither* support tier of Supplementary Figure 10 — neither MAP-robust nor detected in ≥2 studies) and *supported* (at least one axis), and plotted by its per-study-balanced detection frequency (the CRAPome-style promiscuity measure of Supplementary Figure 5C). The fragile mHTT-gains are *far less* promiscuous than the supported gains (median 0.028 vs 0.505, *n* = 21 vs 183, Mann–Whitney^45^ *p* < 10^−6^); the same holds for the losses (median 0.25 vs 0.379). Across both arms, per-call confidence (number of support axes satisfied) and promiscuity are *positively* coupled (Spearman^47^ ρ = 0.795 for gains, 0.799 for losses) — so the fragile calls are *under*-detected, not over-detected, and the “sticky background” explanation is refuted. (The breadth axis and this promiscuity measure are both detection-based, so this coupling is partly definitional; the non-circular control is the inclusion-proteome comparison in Supplementary Figure 8, where inclusion membership does *not* distinguish gains from losses.) Boxes show quartiles and median; whiskers, 1.5×IQR; grey = fragile, coloured = supported.

**Supplementary Figure 8.**
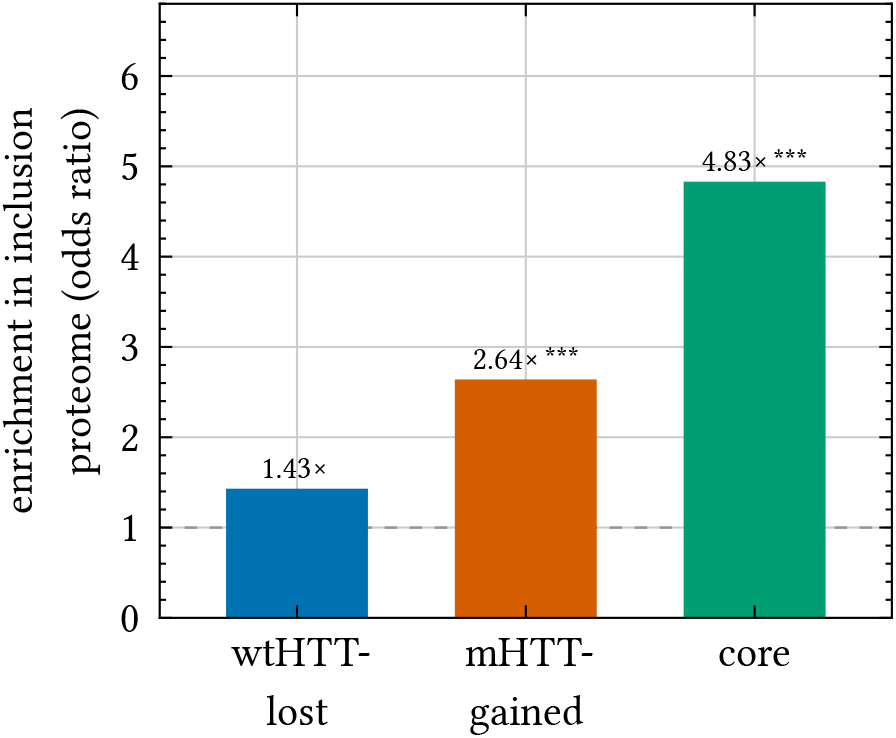
The condition-dependent and -independent HTT interactome is over-represented in the *in vivo* mHTT inclusion proteome. Enrichment (Fisher exact odds ratio^50^ vs the interactome-wide evaluated background; dashed line = no enrichment) of each differential class among proteins sequestered into mHTT inclusions (Hosp *et al*. 2017^33^ — R6/2 insoluble proteome, whole brain, 12 weeks; *n* = 491 of the evaluated proteins). All three classes are significantly over-represented — the constitutive **core** most strongly (4.83×; chaperonins, HSP70 and the proteasome are major inclusion constituents). Notably, inclusion membership does *not* by itself distinguish the mHTT-gains from the wtHTT-losses (gains 2.64× vs losses 1.43×; direct contrast *ns, p* = 0.197): the gain/loss axis rests on the affinity-proteomics evidence (Supplementary Figure 5A), a distinct property from inclusion sequestration. ** *p* < 0.01, *** *p* < 10^−4^.

**Supplementary Figure 9.**
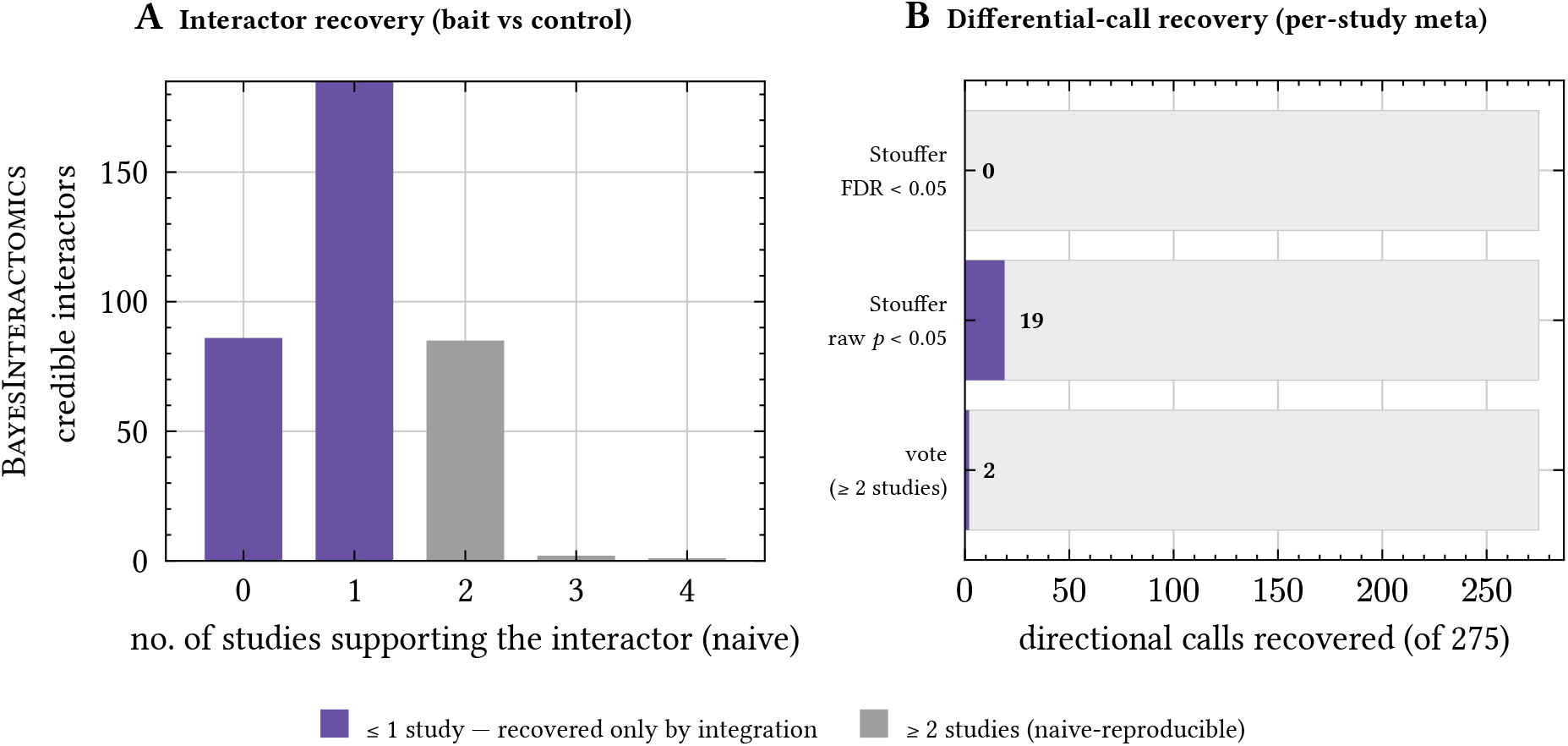
Neither the interactors nor the gained/lost calls are recoverable study-by-study — only the integrated model resolves them. **(A)** BayesInteractomics calls 421 credible interactors (*P* > 0.95 in either bait). Scored *individually* per source study by a conventional rule — detected in ≥50% of that study’s bait immunoprecipitations and ≥2-fold enriched over its controls — each credible interactor is binned by the number of studies that support it. 79% are supported by at most one study, so a naive “reproducible in ≥2 studies” consensus recovers only 88 of the 421 (20.9%). **(B)** The same logic applied to the harder *differential* question. A fair per-study mHTT-versus-wtHTT contrast computed *within* each genotype-split study (Greco, Sap, Justice) on the same MNAR-imputed matrix and combined across studies by sample-size-weighted Stouffer’s Z^43^ — or by direction-consistent vote-counting — recovers 0 of the 275 gained/lost calls at FDR < 0.05^44^ (19 at nominal *p* < 0.05; 2 by vote). The per-study differential signal is individually too weak for any conventional test to call. Because the four datasets overlap little (Supplementary Figure 6), the integrated model borrows strength across them to recover signal — both interactions and their direction — that any single study, or a reproducibility filter, would discard. (The conclusion follows from the low cross-study overlap, not a particular naive threshold.)

**Supplementary Figure 10.**
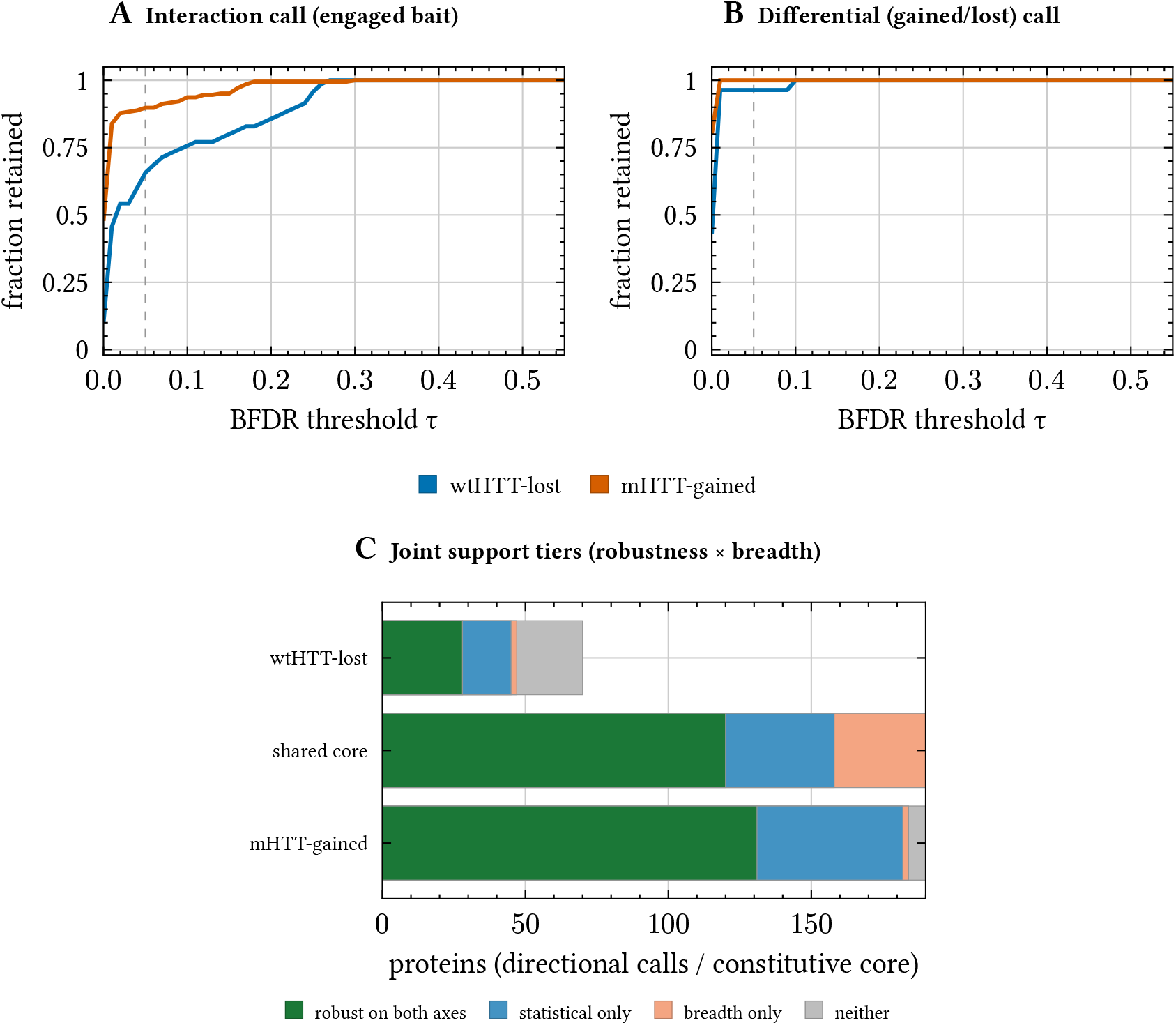
Both the wtHTT-losses and the mHTT-gains are robust interactors and robust differential calls. Each panel plots, for the wtHTT-lost (70 proteins, blue) and mHTT-gained (205 proteins, vermillion) call sets, the *fraction retained* as the Bayesian FDR (BFDR) acceptance threshold τ is tightened from 0.5 towards 0 (x-axis; 1 = whole class survives, 0 = none); the dashed line marks the conventional τ = 0.05. The two panels apply two *different* BFDR criteria to the same proteins. **(A)** *Interaction* robustness: each protein tested against the BFDR for being a genuine interactor *of the bait it engages* (BFDR_wtHTT_ for losses, BFDR_mHTT_ for gains). Both classes hold up — at τ = 0.05, 90% of gains and 66% of losses are retained — so the gains are *bona fide* mHTT interactors, not detection noise. **(B)** *Differential* robustness: each protein instead tested for being *condition-dependent* (gained-vs-unchanged for gains, lost-vs-unchanged for losses). Both classes are well supported on this stricter criterion too: 96% of losses and 100% of gains survive τ = 0.05. That a gained protein binds mHTT, and that it binds mHTT *more than wtHTT*, are therefore both well supported for the majority of gains individually — so we report the gains, like the losses, as per-protein condition-dependent calls. **(C)** Integrating the two axes: each call is binned by whether it is statistically robust (MAP classification agrees with the optimal call) and broadly observed (detected in ≥2 studies, Supplementary Figure 6B), with the constitutive core (unchanged) as the high-confidence anchor. Support forms a gradient — the core is robust on both axes (120/294), and both directional arms retain a sizeable both-axes tier: the wtHTT-losses 28/70 and the larger mHTT-gains 131/205. Statistical robustness and breadth are positively associated within both arms. This two-track assignment is insensitive to where the robustness cut is placed rather than an artefact of a binary threshold: panel B is continuous, with both differential-robustness curves plateauing as *τ* tightens, so sweeping the cut leaves both both-axes tiers essentially unchanged. (The core’s MAP-concordance is partly definitional — the agree-on-*unchanged* set — so breadth is its informative axis.) Of the 275 directional calls, 44 fall in the *neither* tier and rest on the decision-theoretic call alone; reassuringly, no curated functional module is composed mainly of them (at most 29% of any module’s called members), so the functional conclusions are unaffected.

**Supplementary Figure 11.**
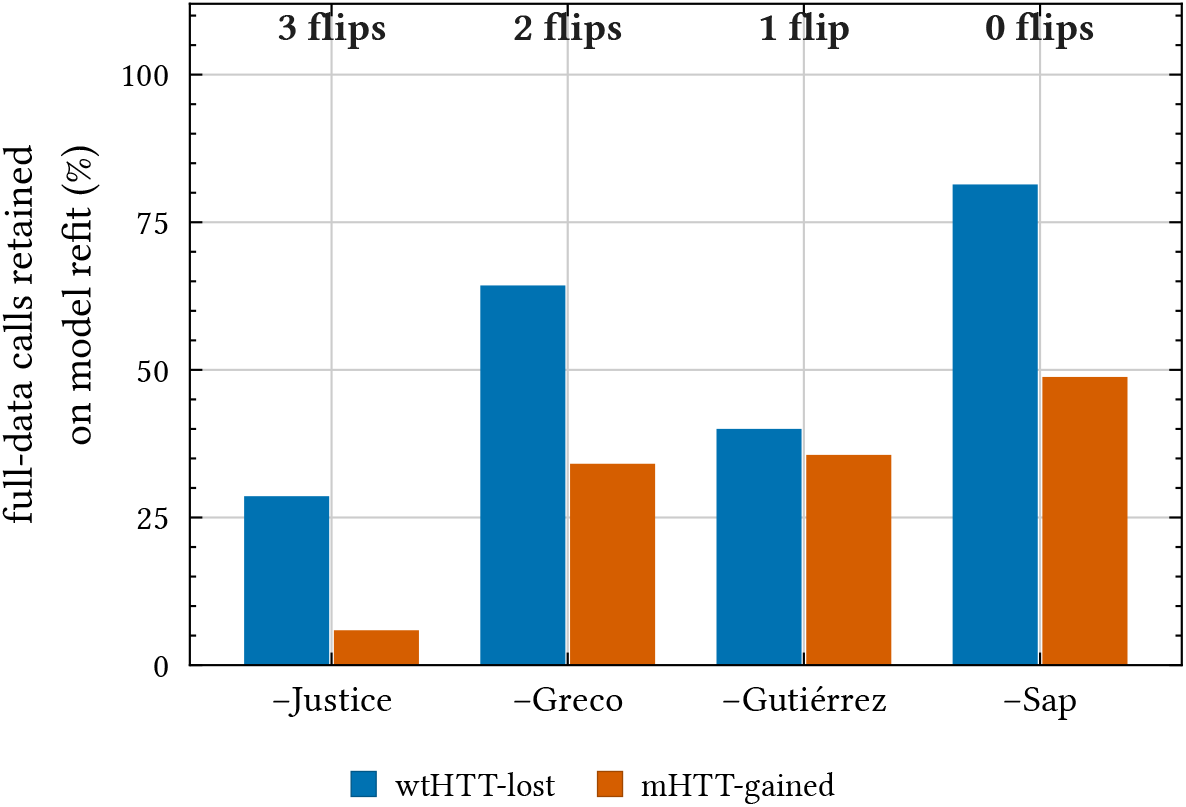
The directional interactome is robust to leave-one-study-out model refits — the residual sensitivity is statistical power, not direction. A true model-refit leave-one-study-out: the entire pipeline (both single-condition models and the wtHTT-versus-mHTT differential) was re-fit four times, each time removing one source study’s protocol(s) from the pooled matrix, and each fold’s calls compared to the full-data calls. Bars give the fraction of the 70 wtHTT-lost (blue) and 205 mHTT-gained (vermillion) full-data calls that keep their direction in each refit; the figure above each fold is the number that *reverse* direction. Across all four refits only 6 of the 275 directional calls ever reverse (97.8% directionally stable; at most 3 in any single fold). Rather than reversing, calls that lose support collapse to non-significant when a study is dropped — mild for the single-protocol datasets, but strong for the dominant Justice2025 (three of six protocols, half the mutant samples; only 6% of gains retained). The mutant arm is the more power-sensitive of the two. This is the model-refit counterpart of the evidence-level leave-one-study-out (Supplementary Figure 6) and quantifies the under-powering of Supplementary Table 3: the pooled model is required because single studies cannot individually resolve the calls.

